# LC-MS/MS characterization of SOBERANA^®^02, a receptor binding domain-tetanus toxoid conjugate vaccine against SARS-CoV-2

**DOI:** 10.1101/2024.12.27.630511

**Authors:** Olivia Martínez, Darielys Santana-Medero, Satomy Pousa, Jean Pierre Soubal, Arielis Rodríguez-Ulloa, Pablo E. Ramos-Bermúdez, Vladimir Besada, Raine Garrido, Paulo Carvalho, Michel Batista, Katharina Zetl, Jacek Wiśniewski, Tamy Boggiano, Yury Valdés-Balbín, Dagmar García-Rivera, Daniel G. Rivera, Vicente Verez-Bencomo, Luis Javier González

**Affiliations:** Finlay Institute of Vaccines, 200 and 21 Street, Havana 11600, Cuba; Center for Genetic Engineering and Biotechnology, Ave 31 e/158 y 190, Havana 10600, Cuba; Carlos Chagas Institute/FioCruz Rua Professor Algacyr Munhoz Mader, 3775 CIC 81350-010, Curitiba, Brazil; Department Proteomics and Signal Transduction, Max Planck Institute of Biochemistry. Am Klopferspitz 18, 82152, Martinsried, Germany; Center for Molecular Immunology, P.O. Box 16040, 216 St., Havana, Cuba; Laboratory of Synthetic and Biomolecular Chemistry, Faculty of Chemistry, University of Havana, Zapata and G, 10400, La Habana, Cuba

**Keywords:** Conjugate vaccines, mass spectrometry, SARS-CoV-2, type 2 peptides, tetanus toxoid, maleimide-thiol chemistry

## Abstract

SOBERANA^®^02 is a safe and effective conjugate vaccine against SARS-CoV-2, produced using the maleimide-thiol chemistry. In this vaccine, the Cys^538^ in the recombinant receptor binding domain (RBD) of SARS-CoV-2 is linked, through a thiosuccinimide linker, to lysine residues of tetanus toxoid (TT) preparation. LC-MS/MS analysis revealed that TT is a complex mixture of proteins, similar to other TTs where the detoxified tetanus neurotoxin (d-TeNT) has been shown to be the most abundant protein (30-56%), regardless the quantification method used. The fifteen most abundant proteins account for approximately 78% of the total proteins. LC-MS/MS analysis of the activation process showed that 102 out of 107 lysine residues in the d-TeNT incorporated a maleimide group. In contrast, when tryptic peptides isolated by Ni^2+^-NTA affinity chromatography, were analyzed by LC-MS/MS, only 22 Lys residues in the d-TeNT were cross-linked to the RBD *C*-terminal tryptic peptide (^538^CVNF^541^-HHHHHH), probably due to steric hindrance. Twelve and eighteen conjugation sites were assigned based on the identification of linear peptides carrying a conjugated lysine residues (Δm = +1454.58 Da or Δm = +1472.59 Da) and cross-linked peptides with stabilized linker forms, respectively. Eight conjugation sites were coincidently assigned by both strategies. The assignment of the conjugation sites was manually validated by observed regularities (z≥3+, XIC, immonium ions, specific linker fragment ions) not considered by the identification software (PEAKS, Kojak and pLink2). The RBD was also conjugated, but to a lesser extent, to ten other low abundance carrier proteins. To our knowledge, this work is the first report of conjugation site assignment in a TT-based conjugate vaccine.

## 1 Introduction

Coronaviruses infections represent a health concern in the 21^st^ century, due to their long-term persistence, enhanced ability of invasion, and the emergence of novel variants of the Severe Acute Respiratory Syndrome Coronavirus 2 (SARS-CoV-2). COVID-19 caused by SARS-CoV-2 virus, is an infectious disease that was firstly originated in Wuhan, China [1]. In January 2020 World Health Organization (WHO) declared COVID-19 as a Public Health Emergency of International Concern. Since the pandemic started, hundreds of millions of people were infected and more than 5 million people lost their lives (https://covid19.who.int). Several technology platforms were used to produce effective COVID-19 vaccines [2] including plasmid DNA vaccines, messenger RNA vaccines, viral vector-based vaccines, bacterial vector-based vaccines, trained immunity-based vaccines and recombinant subunit protein vaccines [3].

To the best of our knowledge, SOBERANA^®^02 is the only protein-protein conjugate vaccine clinically approved and used in vaccination against COVID-19 worldwide. The antigen of SOBERANA^®^02 is a SARS-CoV-2 receptor binding domain (RBD) tetanus toxoid (TT) [4] conjugate vaccine produced by the maleimide-thiol chemistry [5]. An average of six copies of the RBD(R^319^-F^541^)-H_6_ of SARS-CoV-2 expressed in CHO cells, [6] are expected to be linked through the Cys^538^ residue to multiple lysine residues of the carrier protein(s) present in the TT preparation [5]. The six histidine residues (H_6_) were incorporated into the *C*-terminus of the RBD to facilitate the purification process prior to conjugation to TT [6]. SOBERANA^®^02 is administered using a heterologous scheme [7] with two doses of SOBERANA^®^02 followed by a third dose of a SARS-CoV-2 dimeric RBD (SOBERANA^®^ Plus) [6, 8].

In a double-blind, randomized, placebo-controlled phase 3 clinical trial, the vaccine efficacy in this heterologous scheme was 92% against the symptomatic disease [9]. Immunization of children 2-11 years-old with the SOBERANA^®^02/SOBERANA^®^Plus scheme provided strong protection against symptomatic and severe disease against Omicron, a variant of concern. This response was sustained during the six months post-vaccination follow-up study [10].

In our laboratory, accomplishing with the ICH Q6B guidelines [11], we have characterized the recombinant SARS-CoV-2 RBDs produced in different expression systems by using the combination of an in-solution buffer-free digestion method and ESI-MS analysis [12, 13]. This method allowed full-sequence coverage, the identification of the disulfide bonds and the assignment of several post-translational modifications in a single ESI-MS spectrum. This approach was also applied to the characterization of recombinant RBDs as active pharmaceutical ingredient of SOBERANA^®^02 [5], SOBERANA^®^Plus [6] and Abdala, another clinically effective COVID-19 vaccine [14, 15] based on an RBD expressed in *P. pastoris* [16].

The characterization of the SOBERANA^®^02 conjugate vaccine by mass spectrometry, as well as the assignment of the conjugation sites, has not been reported so far. A comprehensive characterization of this vaccine antigen is relevant not only in the context of COVID-19, but also to achieve a better understanding of protein-protein conjugate incorporating the TT carrier protein(s). To our knowledge, there are no previous reports on the characterization by LC-MS/MS analysis of a protein conjugate vaccine antigen based on TT.

The identification of conjugation sites is a quality attribute in conjugate vaccines, because it provides information on the conjugation efficiency and the linkage positions between the antigen and the carrier protein, factors that could modulate the immunogenicity, stability, and safety of the vaccine among other properties [17–19].

To identify the conjugation sites, the homogeneous and heterogeneous conjugate vaccines [20–24] are digested with specific proteases and the complex mixture of linear and cross-linked peptides is analyzed by LC-MS/MS analysis. To identify the conjugation sites, we used the same software [25, 26] that are employed to assign MS/MS spectra to cross-linked peptides in cross-linking mass spectrometry experiments [27]. A particular type of crosslinked peptides, the type 2 peptides [28], are those that contain information of the conjugation sites. A type 2 peptide is composed of two proteolytic peptides, one from the carrier protein and the other from the antigen, which are cross-linked through a spacer resulting from the conjugation technology employed in the vaccine production.

In the case of SOBERANA^®^02 [5], TT is functionalized with multiple maleimide groups by reaction of the lysine (Lys) ε-amino groups with β-maleimidopropionic acid *N*-hydroxysuccinimide ester (BMPS), followed by thiol-maleimide conjugation by Cys^538^ of the recombinant RBD. Therefore, after trypsin digestion, it is expected that all type 2 peptides should be composed of the RBD *C*-terminal tryptic peptide (^538^CVNF^541^-His_6_) linked to different lysine-containing tryptic peptides of the carrier protein(s) by the *N*-propionylthiosuccinimide linker (**Figure 1**, structure (**I**)) and its stabilized forms (**Figure 1**, structures (**III**, **IV** and **V**)).

**Figure 1.**
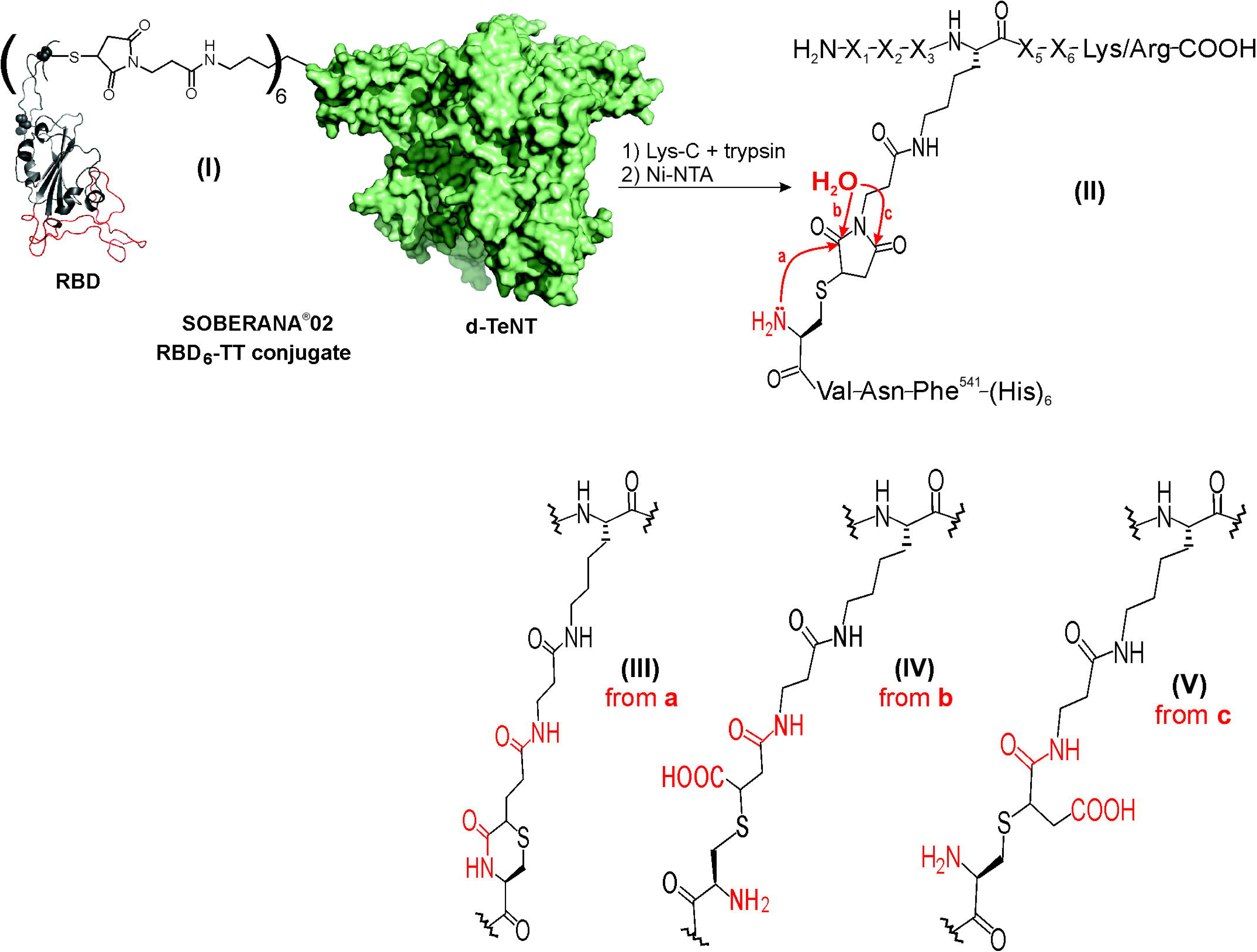
**(a)** Schematic representation of SOBERANA^®^02 (structure (**I**)) synthesized by the maleimide-thiol chemistry as previously reported [5]. This conjugate vaccine is composed by an average of six units of the recombinant RBD of SARS-CoV-2 (Uniprot: P0DTC2, colored in red and black) linked through an *N*-propionylthiosuccinimide linker to the d-TeNT carrier protein (colored in green) of *C. tetani* (Uniparc: UPI000003617E). In the structure of the RBD, the red and black colors indicate the receptor binding motif and the remaining protein structure, respectively. In SOBERANA^®^02, the C^538^ in the RBD antigen is linked to several lysine residues in the d-TeNT carrier protein. Tryptic digestion using a MED-FASP protocol [35] produces a complex mixture of linear and type 2 peptides. The latter were isolated by immobilized Ni^2+^-NTA affinity chromatography and analyzed by LC-MS/MS analysis. Type 2 peptides are composed of the RBD *C*-terminal peptide (^538^CVNF^541^-HHHHHH) and several tryptic peptides of the d-TeNT linked by a thiosuccinimide linker and/or its stabilized linker forms as shown in **(b).** The transcyclization [30, 31] and hydrolysis [32] reactions yield the transcyclized linker (structure (**III**)) and two positional isomers of the hydrolyzed thiosuccinimide linkers (structures (**IV**) and (**V**)), respectively.

Very often, the identification of type 2 peptides is an analytical challenge in the context of a complex proteolytic digest. In heterogeneous conjugates, type 2 peptides are generated by non-quantitative chemical reactions and their ionization efficiency in ESI-MS analysis is lower than the linear peptides. The combination of these two factors is responsible for the low intensity signals of type 2 peptides in LC-MS/MS analysis. Despite this drawback, the use of sensitive mass spectrometers allows the identification of type 2 peptides without requiring their selective isolation or enrichment prior to LC-MS/MS analysis [20–24].

In previous works [22, 29], we have reported that overnight proteolysis or incubation in appropriate conditions (basic pH and 37 °C) converts the original thiosuccinimide linker (structure (**II**), **Figure 1**) of type 2 peptides into the transcyclized (structure (**III**), **Figure 1**) and hydrolyzed (structures (**IV**) and (**V**), **Figure 1**) stable linker forms due to transcyclization [30, 31] and hydrolysis reactions [32, 33], respectively.

The transcyclization involves the nucleophilic attack of the amino group of the *N*-terminal cysteine to the carbonyl group in the succinimide ring (red curved arrow, structure (**II**), **Figure 1**). As result of this internal rearrangement, the original five-membered ring thiosuccinimide linker is transformed into a transcyclized linker composed by a six-membered ring (structure (**III**), **Figure 1**). There are two requisites for this transcyclization to take place, the Cys involved in the thioether bond should be placed at the *N*-terminal end and its amino group should free and non-protonated. On the other hand, the imide bond in the thiosuccinimide linker can be hydrolyzed by two pathways (pathways (b) and (c), structure (**II**), **Figure 1**) to yield two positional isomers of the thiosuccinamic acid thioether (structures (**IV**) and (**V**), **Figure 1**).

Considering that all type 2 peptides may have this structural heterogeneity in the linker and being aware that they provide valuable information about the conjugation sites, both linker forms need to be included in the analysis of conjugate vaccines [20, 22, 24]. For the identification of the conjugation sites of SOBERANA^®^02, we anticipated additional challenges due to the number of potential carrier proteins present in the TT preparation. In 2019, Möller *et al* [34] characterized by LC-MS/MS analysis the protein composition of TT-based vaccines from five different manufacturers. They reported that the formaldehyde detoxified tetanus neurotoxin (d-TeNT) was the most abundant protein, ranging from 30% to 60%, depending on the manufacturer and the quantification method used [35–37]. LC-MS/MS analysis of the tryptic digests revealed that individual tetanus vaccines are a very complex mixture of 360 to 670 proteins. A total of 991 proteins were identified in this study. All preparations shared a common core of 206 proteins containing 54 potential antigens as predicted by the Vaxijen software [38]. These results imply, that the complexity of the peptide mixture generated by proteolysis would be higher than that of generated by conjugate vaccines produced and characterized earlier in our group from a single pure carrier protein [20–24].

Additionally, in the TT production process, formaldehyde introduces chemical modifications such as Schiff base (+12.000 Da), the methylol (+30.011 Da) and crosslinks of several amino acids (K, R, Q, N, W, and Y), [39, 40] in the proteins secreted by *C. tetani*. All these aspects should be taken into account for the mass spectrometric characterization of SOBERANA^®^02, and they suggest that the application of proteomics tools is mandatory.

The three main goals of our work were: (1) to report the protein composition of the TT preparation used in SOBERANA^®^02; (2) to study the activation process of TT for further conjugation and, (3) to identify the conjugation sites in SOBERANA^®^02.

## 2 Materials and Methods

The tetanus toxoid preparation obtained from *C. tetani* (strain Harvard) (batch EC-TTC-2105) was produced at the Finlay Institute of Vaccines in Havana, Cuba. The recombinant receptor binding domain of SARS-CoV-2 (RBD (R^319^-F^541^)-H_6_) expressed in Chinese hamster ovary (CHO)-K1 cells (batch 5028/P2112) was provided by the Center for Molecular Immunology [6] in Havana, Cuba. The conjugate vaccine SOBERANA^®^02 (batch 1021TS2) was synthesized by the Finlay Institute of Vaccines as described before [5]. Other solvents and reagents were purchased from commercial suppliers.

### 2.1 Activation process of TT preparation

A solution of TT (10 mL, 5 mg.mL^−1^, 50 mg, MW 150 kDa, 0.33 μmol) in 100 mM 4-(2-hydroxyethyl)piperazine-1-ethanesulfonic acid (HEPES) pH 7.8 reacted with a freshly prepared solution of BMPS (3-maleimidopropionic acid *N*-hydroxysuccinimide (NHS) ester) in DMSO (187 μL, 75 mg.mL^−1^, 53 μmol, 160 equivalent per mol of TT). The mixture was gently stirred for 1 h at room temperature, purified by diafiltration (100 kDa cut-off) with 35 mM PBS pH 7.4, 5 mM EDTA, and finally concentrated to 18 mg/mL. The stoichiometry of maleimide incorporated to TT was found to be between 20 - 30 maleimide groups per TT, as measured by a reversed Ellman method [41]. To block the maleimide groups in the carrier protein(s), 75 equivalents of cysteamine hydrochloride (Tokyo Chemical Industry, Japan) were added per mole of TT and the reaction was stirred for 30 min at 2 - 8 °C under nitrogen atmosphere (see **Supplemental file 1**, **Fig. S1**). The resulting solution was diafiltrated with 1 mM, pH 7, PBS buffer solution. The reaction completion was verified by the reverse Ellman’s method [41].

### 2.2 Proteolytic Digestions

All proteolytic digestions of the TT preparation, activated TT preparation and SOBERANA^®^02 conjugate vaccine proceeded according to the MED-FASP protocol [35] with slight variations. A volume containing 0.2 mg of the protein mixture dissolved in 50 mM DTT and 1.3% SDS was kept at 95 °C for 5 min. The solution was cooled down to room temperature and then *S*-alkylated with a iodoacetamide solution at a final concentration of 125 µM in the dark for 30 min.

Then, the mixture of the reduced and *S*-alkylated proteins was filtered through a 30 kDa unit (Microcon-30, Millipore, USA). Samples were later washed with 2 mL of urea buffer (8 M Urea, 100 mM, pH 8.5, Tris-HCl) to remove SDS. The filter device was washed with 600 µL of digestion buffer (50 mM, pH 8.5, Tris-HCl). Samples were incubated with 2 µg of Lys-C (WAKO, Japan), for 16 h at 37 °C. Each digestion was filtered at 4000xG and collected in different tubes. Protein mixture was digested with 1.5 µg trypsin (SIGMA) at 37 °C and chymotrypsin (SIGMA) during 8 h each, and the resultant proteolytic peptides were collected following the same procedure. For the identification of the conjugation sites, only the tryptic digestion was analyzed by LC-MS/MS. Peptides mixture was desalted using Stage tips procedure following manufacturer instructions (ThermoScientific, USA). The peptide concentration was determined for each filtrate using the tryptophan fluorescence [42].

### 2.3 Enrichment of type 2 tryptic peptides derived from SOBERANA^®^02 by using Ni^2+^-NTA affinity chromatography

Ni^2+^-NTA affinity matrix (Pharmacia, Sweden) was equilibrated in 100 mM NH_4_HCO_3_ buffer (pH = 8.3) at 4 °C. Tryptic peptides dissolved in the same equilibrium buffer were mixed with the equilibrated matrix and incubated for 2 h at 4 °C with a gentle shaking. The type 2 peptides containing the *C*-terminal peptide of the RBD (^538^CVNF^541^-H_6_) were retained on to the Ni^2+^-NTA affinity matrix. The column was washed with the same equilibration buffer to remove mainly the contaminant linear and other formaldehyde cross-linked peptides. One percent TFA solution was added until pH = 2.0 and the mixture was centrifuged at 14 000 rpm. This process was repeated twice. The eluted peptides were desalted using a C18 Secpack cartridge (Waters, USA) and eluted in an acetonitrile solution containing 0.2% formic acid, dried in the speedvac, resuspended in 1% formic acid solution and were freeze at -20 °C until LC-MS/MS analysis.

### 2.4 LC-MS/MS analysis

LC-MS/MS analyses were performed at the mass spectrometry facility RPT02H/Carlos Chagas Institute–Fiocruz Paraná. The runs were performed on an Ultimate 3000 nLC (Thermo Scientific coupled with an Orbitrap Fusion Lumos Mass Spectrometer (Thermo Scientific). The peptide mixture was resuspended in formic acid 0.1% and quantified by absorbance in nanodrop (Thermo Scientific). Then, 0.5 µg was loaded on a 15 cm length, 75 µm I.D. emitter, packed in-house with ReproSil-Pur C18-AQ 3 µm resin (Dr Maisch GmbH HPLC) for peptide separation. The column was equilibrated with 250 nL/min of 95% of mobile phase A (0.1% formic acid in water) and the mobile phase B (0.1% formic acid, 5% water in acetonitrile) was increased linearly up to 40% in 60 min. The instrument was set to data-dependent acquisition mode with time between master scans set to 1 s and exclusion list of 15 s. Survey scans (350 – 2000 m/z) were acquired in the Orbitrap system with a resolution of 60,000 at m/z 200. The most-intense ions with charge states from +2 to +7 were sequentially isolated and fragmented in the stepped higher-energy collisional dissociation (HCD) collision cell using normalized collision energies of 27, 30 and 33. The fragment ions were analyzed with a resolution of 15 000 at 200 *m/z*. The general mass spectrometric conditions were as follows: 2.30 kV spray voltage, capillary temperature of 175 °C, predictive automatic gain control (AGC) set to 250% on MS1 and 220% for MS2.

### 2.5 Protein identification by LC-MS/MS

The TT preparation was digested with Lys-C, trypsin and chymotrypsin using the MED-FASP method [35]. The three filtrates of these protein digests were separately analyzed by LC-MS/MS, and the raw files were loaded in the MaxQuant (v2.0.3.1) [43] and PEAKS Studio V7.0 [44] for protein identification. The database search was carried out on the *C. tetani* (strain Harvard) proteome reported in the UNIPARC database (UP000290279) containing 2708 entries (downloaded 13^th^ June 2024, https://www.uniprot.org). Only one missed cleavage site was allowed for the identified peptides. Proteins with at least one unique peptide and a false discovery rate of 1% was allowed in the final output. The mass tolerance for the matching peptides were 10 ppm. The correspondence between proteins in the *C. tetani* proteomes available in the UNIPARC (strain Harvard, UP000290279) and UNIPROT (strain Massachusetts, UP000290921) was carried out with the ID mapping tool available in the UNIPROT website. Absolute quantification protein abundance indexes (TOP3 [37], iBAQ [36] and TPA [35]) were calculated by the MaxQuant software (v2.0.3.1) [43].

### 2.6 Bioinformatic analysis of top fifteen most-abundant proteins identified in the TT preparation from C. tetani (strain Harvard)

Biological functions associated with top fifteen most-abundant proteins of *C. tetani* identified in the TT preparation were extracted from the UniProtKB database (https://www.uniprot.org/ Accessed on January 2023) [45]. Proteins cellular localization was predicted by PSORTb v3.0.3 [46]. Prediction of the putative antigen function of proteins was performed using the VaxiJen database (http://www.ddg-pharmfac.net/vaxijen/VaxiJen/VaxiJen.html, accessed on January 2023) with a threshold of 0.4 [38]. To find similarity with other proteins sequences, proteins without functional annotations were analyzed with BLASTp (https://blast.ncbi.nlm.nih.gov/Blast.cgi?PAGE=Proteins, accessed on April 2024) [47]. The search was performed using as standard database the UniProtKB/Swiss-Prot, other parameters were setting in its default values.

### 2.7 Identification of the conjugation sites in SOBERANA^®^02 by LC-MS/MS analysis

The identification of the conjugation sites was based on two strategies. In the first strategy, the MS/MS spectra were assigned to linearized peptides containing modified lysine residues. The molecular masses of the modified lysine residues were increased either by 1454.58 Da or 1472.59 Da when linked to the *C*-terminal tryptic peptide of the RBD via a transcyclized or a hydrolyzed thiosuccinimide linker, respectively. The remaining database search parameters for protein identification using PEAKS Studio V7.0 [44] were the same as described in section 2.5.

The second strategy was based on the assignment of MS/MS spectra to type 2 tryptic peptides by using the pLink2 (v2.3.11) [25] and Kojak (v1.6.0) [26] software. The presence of the stabilized forms of the thiosuccinimide linker (hydrolyzed and transcyclized forms), were taken into account for the identification of type 2 tryptic peptides. The elemental compositions of the transcyclized and the hydrolyzed thiosuccinimide linkers were C_7_H_5_O_3_N and C_7_H_7_O_4_N, respectively. The cross-linked amino acids for all linker forms were the primary amino groups (lysine and the *N*-terminal end of the proteins) and the cysteine residues. The allowed precursor and fragment ion mass tolerances were 10 and 15 ppm, respectively. The minimum and maximum molecular mass limits for the identified type 2 peptides were 400 and 8000 Da, respectively. Up to two missed cleavage sites were allowed for trypsin digestion. Deamidation of N/Q, *S*-carbamidomethylation of Cys, and the oxidation of Met residues were defined as variable modifications. The database contained the sequences of the recombinant RBD(^319^R-F^541^)-H_6_ from SARS-CoV-2 [6] and the fifteen most abundant proteins in the TT preparation. The database used with Kojak software additionally contains all the reverse sequences to determine the false discovery rate (FDR) through the Percolator (v2.08) software [48]. A 5% of FDR was allowed in the software output. Other search parameters were the same as previously reported [20]. The set of MS/MS spectra assigned to type 2 peptides was manually validated by considering the sequence coverage of both cross-linked peptides in the MS/MS spectra. The linker fragment ions generated by the dissociation of the newly formed pseudopeptide in the transcyclized (P+71 and C+80 ions, **Supplemental file 1**, **Fig. S2**) [29] and hydrolyzed (P+71 and C+98 ions, **Supplemental file 1**, **Fig. S3**) linkers [20, 49] were considered in the output data validation. With the same goal, the thioether bond fragmentation (P+169, P+203, C-SH and C-34 ions, [23, 29] and other linker fragment ions were also taken into consideration (**Supplemental file 1**, **Fig. S4 -S6**). Moreover, in the manual validation, we also considered other backbone fragment ions derived from the two component linear peptides resulting from the linker fragmentation carrying modified amino acids (Lys and Cys).

## 3 Results and Discussion

### 3.1 Protein composition of the TT preparation used as the carrier protein in SOBERANA^®^02

As a first step in this research, the protein composition of the TT preparation used to produce SOBERANA^®^02 was determined by MED-FASP digestion protocol [35] and LC-MS/MS analysis. The protein sequence database of *C. tetani* (strain Harvard, 2708 entries) was searched using PEAKS [44] and MaxQuant [43] software. By overlapping the results of both software, 909 different proteins were identified (**Supplemental file 2**). PEAKS and MaxQuant software identified 777 and 695 proteins, respectively (**Supplemental file 1**, **Fig. S7-S9)**. The Venn diagram shows that a common core of 563 proteins, representing approximately 62% of the total proteins, were identified by both software (**Supplemental file 1**, **Fig. S7**). The software PEAKS [44] and MaxQuant [43] exclusively identified 214 and 132 different proteins, respectively (**Supplemental file 1**, **Fig. S7** and **Supplemental file 2**). The results indicate that the TT preparation used in SOBERANA^®^02 has a complex protein composition, similar to other five commercial vaccines against *C. tetani* previously characterized by LC-MS/MS according to Möeller *et al* [34].

In this preparation, the formaldehyde detoxified tetanus neurotoxin (d-TeNT) was the most abundant protein regardless the quantification method used [35–37] (**Table 1**). The d-TeNT abundance ranged from 32.9% to 54.5% when TOP3 [37] and TPA [35] quantification methods were used, respectively (**Table 1**). The second most abundant protein (UPI0002DAC767, Putative S-layer protein/N-acetylmuramoyl-L-alanine amidase) represents approximately 10% of the total proteins while the remaining proteins have abundances below 5% (**Table 1**).

**Table 1.** The top 15 abundant *C. tetani* proteins identified by LC-MS/MS analysis of the tryptic digests of the tetanus toxoid preparation used for the synthesis of SOBERANA^®^ 02.

| #<sup>a)</sup> | UniPARC/<br>Uniprot entries<br>(MW, # K residues) | Annotation | Function | Localization<sup>b)/</sup><br>Probable<br>Antigen<sup>c)</sup> | Abundance<br>(average)/<br>iBAQ/TPA/<br>TOP3/<sup>d)</sup> |
| --- | --- | --- | --- | --- | --- |
| <u>1</u> | UPI000003617E/<br>P04958<br>(151 kDa, 107 Lys) | Tetanus<br>neurotoxin * | Inhibits<br>neurotransmitter<br>release | S / Yes | <u>(44.9)</u>/47.6/<br>54.5/ 32.9/ |
| <u>2</u> | UPI0002DAC767/<br>Q898I7<br>(131 kDa, 168 Lys) | Putative S-layer<br>protein/N-acetylmuramoyl-L-alanine<br>amidase | - | CW / Yes | <u>(10.2)</u>/13.0/<br>9.9/7.6/ |
| <u>3</u> | UPI00001A9618/Q8<br>97C2<br>(52 kDa, 30 Lys) | Tyrosine phenol-lyase * | Tyrosine metabolic<br>process | C / Yes | <u>(4.8)</u>/5.2/<br>2.8/6.3/ |
| <u>4</u> | UPI00000106CD/Q<br>891F6<br>(31 kDa, 22 Lys) | 3-hydroxy-<br>butyryl-CoA<br>dehydrogenase<br>* | Butanoate<br>metabolism | C / No | <u>(4.7)</u>/4.2/<br>4.8/ 5.1/ |
| <u>5</u> | UPI00016089BF /<br>A0A4Q0VB59<br>(58 kDa, 46 Lys) | Chaperonin<br>GroEL,<br>Chaperonin-60,<br>Cpn60* | Protein folding.<br>The GroEL-GroES<br>system promotes<br>and accelerates<br>protein folding | C / Yes | <u>(2.7)</u>/3.3/<br>2.0/2.7/ |
| <u>6</u> | UPI0000010752 /<br>Q890S3<br>(46 kDa, 27 Lys) | Methyl-aspartate<br>ammonia-lyase * | Amino-acid<br>degradation | C / Yes | <u>(2.6)</u>/2.6/<br>1.9/3.4/ |
| 7 | UPI000001003C/Q8<br>97Y7<br>(45 kDa, 39 Lys) | Predicted<br>Amidohydrolase<br>* | Hydrolase activity | C / Yes | <u>(1.5)</u>/1.4/<br>1.4/1.6/ |
| 8 | UPI000000FF03/<br>Q899E8<br>(14 kDa, 11 Lys) | Translation<br>initiation inhibitor<br>* | Organonitrogen<br>compound<br>catabolic process | C / No | **<u>(1.06)</u>**/0.8/<br>1.0/1.4/ |
| **<u>9</u>** | UPI000001053D/<br>Q892V6<br>(80 kDa, 86 Lys) | Cell wall-binding<br>repeat-containing<br>protein * | - | CW / Yes | **<u>(1.05)</u>**/1.0/<br>0.9/1.2/ |
| **<u>10</u>** | UPI0002DA4B64/<br>Q897Y8<br>(44 kDa, 38 Lys) | Predicted<br>amidohydrolase<br>* | Hydrolase activity | C / Yes | **<u>(0.98)</u>**/0.9/<br>0.8/1.1/ |
| **<u>11</u>** | UPI000001030F/<br>Q895B4<br>(97 kDa, 82 Lys) | Aldehyde-<br>alcohol<br>dehydrogenase<br>* | Carbon utilization;<br>Alcohol metabolic<br>process | C / Yes | **<u>(0.9)</u>**/0.91/<br>1.03/0.86/ |
| **<u>12</u>** | UPI000001056B/<br>Q892R0<br>(66 kDa, 51 Lys) | Chaperone<br>protein DnaK,<br>HSP70 * | Acts as a<br>chaperone. Stress<br>response. | C / Yes | **<u>(0.89)</u>**/0.62/<br>0.5/1.5/ |
| 13 | UPI000000FF89/<br>Q898R4<br>(36 kDa, 29 Lys) | Glyceraldehyde-3-phosphate<br>dehydrogenase | Carbohydrate<br>degradation,<br>glycolysis | C / Yes | **<u>(0.76)</u>**/0.75/<br>0.72/0.82/ |
| **<u>14</u>** | UPI000001011B/<br>Q897B4<br>(20 kDa, 17 Lys) | Protein with<br>rubredoxin/<br>rubrerythrin<br>domain* | Electron transport | C / Yes | **<u>(0.67)</u>**/0.47/<br>0.39/1.14/ |
| 15 | UPI0000010662/<br>Q891R3<br>(60 kDa, 54 Lys) | Formate-tetrahydrofolate<br>ligase * | One-carbon<br>metabolism | C / Yes | **<u>(0.41)</u>**/0.35/<br>0.37/0.52/ |

a) Bold and underlined numbers indicate those proteins that were found conjugated to the RBD of SARS-CoV-2 (see Table 2-4 and Supplemental Tables S1-S3).
b) Subcellular localization predicted with PSORTb v.3.0.3 (S: secreted, CW: cell wall, C: cytoplasm).
c) The prediction of probable antigen proteins was performed with VaxiJen [38] bioinformatic server (http://www.ddg-pharmfac.net/vaxijen/VaxiJen/VaxiJen.html).
d) The values in this column indicate the decreasing order in the abundance of the top 15 abundant proteins determined in the LC-MS/MS analysis of the tryptic digestion. The abundance was determined by averaging the values provided by three different quantification methods, iBAQ [36], TPA [35] and TOP3 [37]. The values indicated in bold and underlined correspond to the average of the three quantification methods used.
\* Indicates those proteins identified in other five toxoid preparation of commercially available Tetanus vaccines, according to results reported by Möller J *et. al.* [34].

Based on these results, we constructed a sub-database with the fifteen most abundant potential carrier proteins aiming to identify the potential conjugation sites in SOBERANA^®^02 (**Table 1**). For the ranking list, we considered an average of the abundance values provided by the three individual quantification methods used [35–37]. The abundance of the protein ranked fifteenth in this list (**Table 1**) is lower than 0.5%. We also considered the identification of conjugation sites in carrier proteins with lower abundance to be unlikely.

### 3.2 Bioinformatic analysis of the fifteen most-abundant carrier proteins present in the analyzed TT preparation

LC-MS/MS analysis of five TT-based vaccines, demonstrated that besides d-TeNT (UPI000003617E), other proteins of *C. tetani* are also present in toxoid preparations [34]. In fact, 13 out of the fifteen most abundant proteins that were identified in our study have been previously identified in five commercially available TT preparations (see asterisks in Annotation of **Table 1**) [34]. In this TT preparation thirteen proteins, others than d-TeNT, were predicted as antigens based on VaxiJen [38] database screening (**Table 1**). Of note, two potential antigens were predicted to be localized in the extracellular space as part of the *C. tetani* cell wall. Among them, the protein ranked second in abundance (UPI0002DAC767) is a structural component of the S-layer, such structure allows bacteria to adhere to surfaces and maintain shape and envelope rigidity. Besides the cell wall-binding repeat-containing protein (UPI000001053D) has similarity to the *Bacillus subtilis* N-acetylmuramoyl-L-alanine amidase LytC (**Table 1**). Such cell wall remodeling enzymes plays an essential role in mediating contact-dependent exchange of molecules among bacterial cells [50]. Hypothetically all these proteins may contribute to protection against *C. tetani* infection.

Among the fifteen most-abundant proteins, cytoplasmic localization has also been predicted (**Table 1**). Cell lysis during the TT production might explain the accumulation of highly abundant and stable cytoplasmic proteins of *C. tetani*. For instance, proteins related to amino acids degradation (Methylaspartate ammonia-lyase, protein # 6) and metabolism (Tyrosine phenol-lyase, protein # 3), protein folding (Chaperonin GroEL, protein # 5), carbon metabolism (Glyceraldehyde-3-phosphate dehydrogenase, protein # 13, and Formate-tetrahydrofolate ligase, protein # 15) were identified (see **Table 1**).

### 3.3 Activation process

The BMPS-activated TT preparation (**Supplemental file 1**, **Fig. S1**) was digested with Lys-C, trypsin and chymotrypsin by MED-FASP protocol [35] and the filtrates were independently analyzed by LC-MS/MS. In d-TeNT (UPI000003617E), we found that 86 out of the 107 lysine residues (**Supplemental file 1**, **Fig. S10-S11** and **Supplemental file 3**) were partially acylated at their epsilon amino group with either the intact *N*-propionylmaleimide group (K(+151.03), in total 22 Lys) or its hydrolyzed form (K(+169.04), in total 64 Lys), (**Supplemental file 1**, **Fig. S10-S11** and **Supplemental file 3**). The latter being the most frequently observed was favorably generated due to the basic pH during proteolysis [33]. This result is consistent with a non-specific activation process [5], which targets basic and hydrophilic lysine residues generally exposed on the protein surface. We also found these lysine residues in their unmodified variant confirming that this activation process with BMPS proceeded partially. These results demonstrate that d-TeNT protein (UPI000003617E) has multiple potential conjugation sites that can be linked through a *N*-thiosuccinimidopropionyl linker to the RBD Cys^538^ (**Fig. 1**). We also found evidence of activation of lysine residues in other proteins of *C. tetani* present in the tetanus toxoid preparation (data not shown).

When the BMPS-activated TT preparation was reacted with cysteamine (**Supplemental file 1**, **Fig. S1**); digested by the MED-FASP protocol; and analyzed by LC-MS/MS, we found that cysteamine (used as a model for a thiol group donor), was linked to 83 and 102 lysine residues through an intact (K(+228.06) and hydrolyzed (K(+246.07)) *N*-propionylthiosuccinimide linker, respectively (**Supplemental file 1**, **Fig. S12**-**S14** and **Supplemental file 4**).

The results demonstrate that a considerable number of lysine residues (102 out of 107) are activated with the BMPS and therefore, become are available for the addition of thiol-containing molecules. However, cysteamine is a small molecule that can reach nearly all incorporated maleimide groups due to the lack of steric hindrance, so this result might not fully reproduce the reaction between BMPS-activated TT and a recombinant protein of around 30 KDa like the RBD.

### 3.4 Identification of conjugation sites in SOBERANA^®^02 by LC-MS/MS analysis

In a first attempt to identify the conjugation sites in SOBERANA^®^02, we applied the same strategy as previously used in our laboratory [20, 21] for the characterization of other heterogeneous conjugate vaccines in which several copies of a peptide or a protein are conjugated a carrier protein. This strategy is based on the LC-MS/MS analysis of the conjugates digested with specific proteases, the assignment of the MS/MS spectra to type 2 peptides using dedicated software [25, 26], and data validation [22, 23].

Unfortunately, this strategy was unsuccessful despite the use of a very efficient protein digestion protocol [35] and a modern mass spectrometer covering a wide dynamic concentration range. Probably the proteolytic digestion of such a complex mixture of carrier proteins (more than 900, see **Supplemental file 2** and **Supplemental file 1, Fig. S7**) in the TT preparation could be a reason for this negative result. Proteolytic digestion significantly increases the complexity of the sample analyzed by LC-MS/MS. Linear peptides are ionized more efficiently than type 2 peptides, and the latter are generated by non-quantitative chemical reactions. The combination of these two factors makes difficult the automatic selection of type 2 peptides for MS/MS analysis.

However, type 2 peptides derived from SOBERANA^®^02 have in their structures the six histidine residues incorporated at the RBD *C*-terminus (**Figure 1a**). Therefore, they could be enriched prior to LC-MS/MS analysis in the same way as the RBD was purified by IMAC affinity chromatography [6].

In this new attempt, when a Ni^2+^-NTA chromatography step was included before LC-MS/MS analysis of the proteolytic digest, thirty-six MS/MS spectra were assigned to eighteen type 2 tryptic peptides with transcyclized linker (structure (**III**), **Figure 1**) by overlapping the outputs of pLink2 [25] and Kojak [26] software (**Table 2** and **Supplemental Files 5-6**). In all of them, the C^538^ in the *C*-terminal peptide of the RBD (^538^CVNF^541^-H_6_) is cross-linked to several lysine residues of tryptic peptides derived from d-TeNT of *C*. *tetani* (**Table 2**).

**Table 2.**
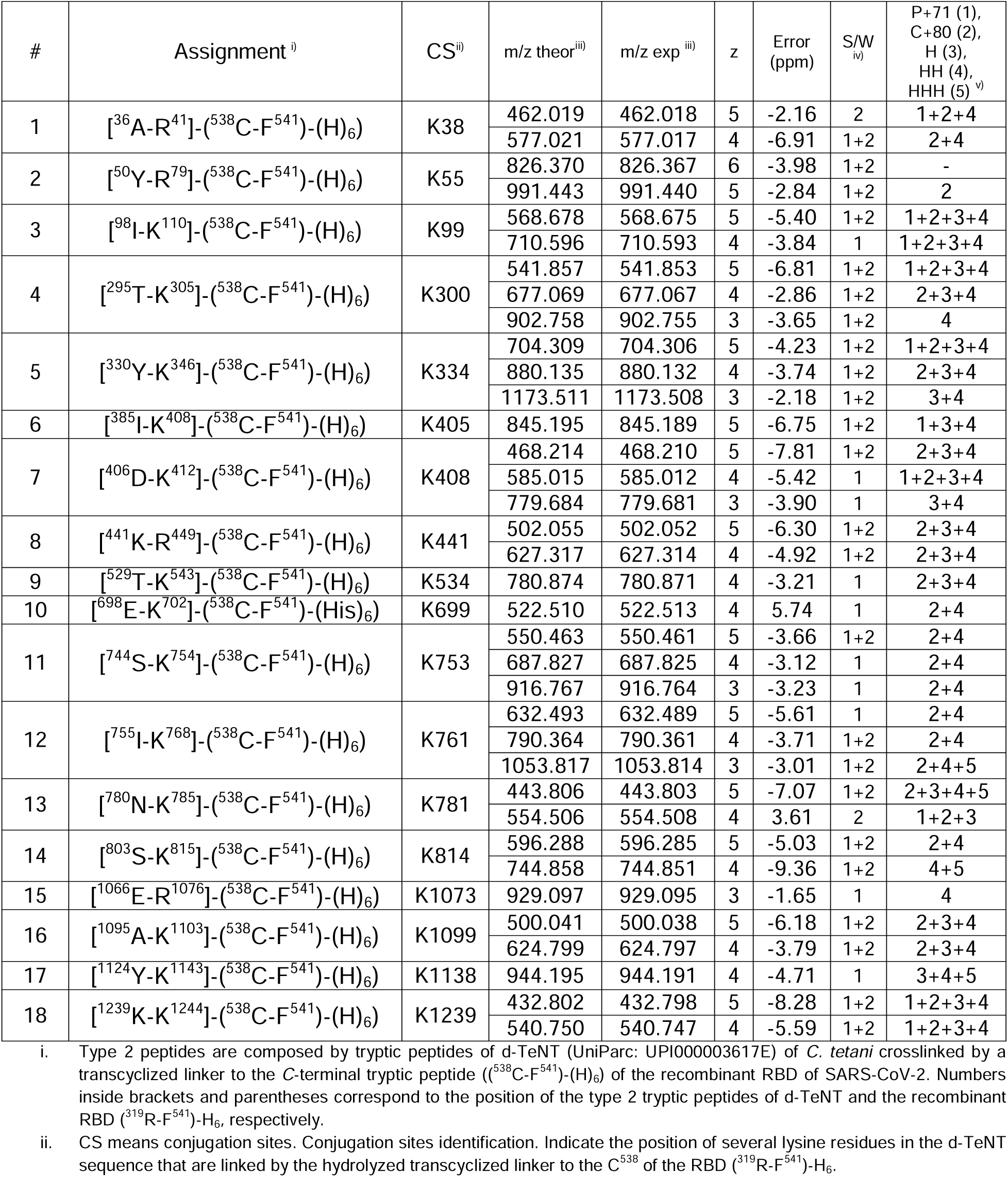
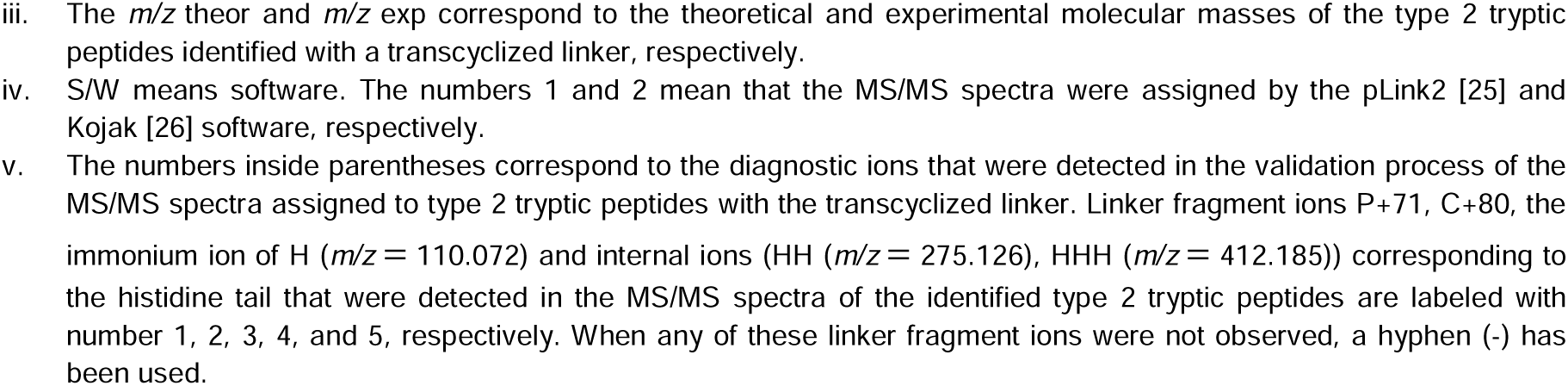
Conjugation sites identification in the d-TeNT (UPI000003617E) determined by LC-MS/MS analysis and based on the assignment of type 2 peptides with a transcyclized linker.

**Figure 2** shows the MS/MS spectrum assigned to a type 2 tryptic peptide ([^1239^KMEAVK^1244^]-(^538^CVNF^541^-H_6_)) with the transcyclized linker. The linker fragment ions P+71 (*m/z* = 776.4327, 1+ and *m/z* = 338.720, 2+) and C+80 (*m/z* = 692.779, 2+ and *m/z* = 462.187, 3+), generated by the fragmentation of the newly formed pseudopeptide bond in the transcyclization reaction, were detected [22].

**Figure 2.**
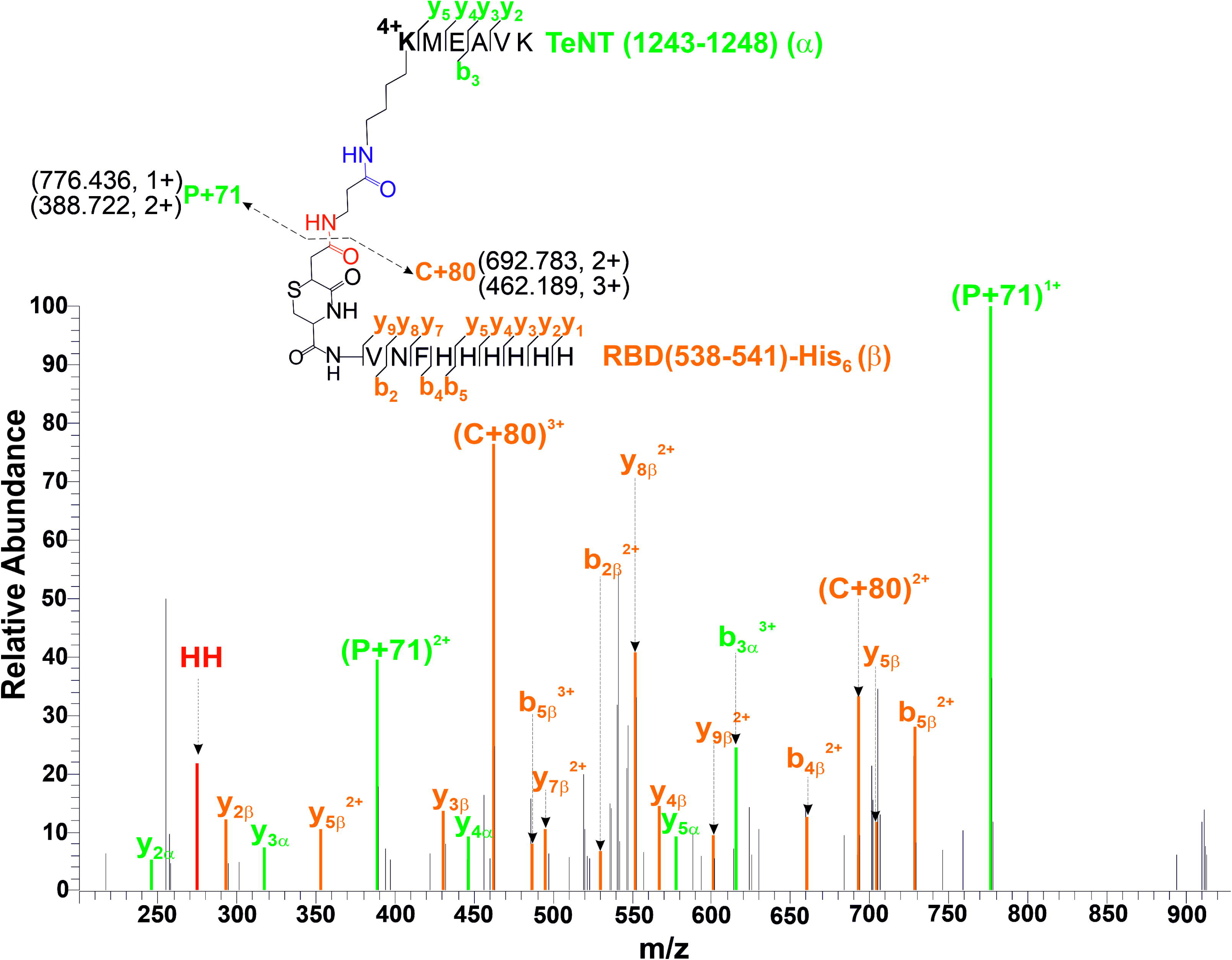
ESI-MS/MS spectrum assigned by pLink2 software [25] to a tryptic type 2 [^1239^**<u>K</u>**MEAVK^1244^]-(^538^**<u>C</u>**VNF^541^-H_6_) is composed by two peptides, one derived from the RBD of SARS-CoV-2 (^538^**<u>C</u>**VNF^541^-H_6_) and the other [^1243^**<u>K</u>**MEAVK^1248^] from the d-TeNT (UPI000003617E) of *C. tetani*. C^538^ and K^1239^ in the RBD of SARS-CoV-2 and d-TeNT from *C. tetani* are crosslinked *via* a transcyclized linker. The (P+71) and (C+80) linker fragment ions detected at (*m/z* = 776.433, 1+ and *m/z* = 338.720, 2+) and (*m/z* = 692.779, 2+ and *m/z* = 462.187, 3+), correspond to the fragmentation of the pseudopeptide bond [22] newly formed due to the transcyclization reaction (highlighted in red in Figure 1, structure (**III**)) [30, 31]. The signals at *m/z* = 110.070, 1+ and *m/z* = 275.126, 1+ correspond to the immonium ion of the histidine (H) and an internal fragment ion (HH) of the tandem repeat of six histidine residues present in type 2 tryptic peptides identified here, respectively.

Although type 2 peptides with the thiosuccinimide and the transcyclized linkers are isobaric species, it seems that most of them were detected with the transcyclized linker form. In 29 out of 36 MS/MS spectra either the C+80 or P+71 linker fragment ions or both were observed in the MS/MS assigned to type 2 peptides (**Table 2**). These linker fragment ions are only permissive for the transcyclized linker [29]. The linker fragment ion most frequently observed in the MS/MS spectra was the C+80 (in 28 out of 36 MS/MS spectra). It suggests that the six-histidine tail in the C-terminal tryptic peptide is very efficient in retaining protonation in gas phase after linker fragmentation and thus facilitating its detection in the MS/MS spectra.

The transcyclization reaction [30] is not possible for this conjugate vaccine because C^538^, which is involved in chemical conjugation, is an internal amino acid residue. However, the trypsin cleavage site at K^537^ (**Figure 1**) generates the *C*-terminal peptide of the RBD (^538^CVNF^541^-H_6_) with C^538^ at the *N*-terminal end (**Figure 1a**). This sequence feature in the RBD sequence combined with the proteolysis conditions (basic pH, 37 °C for 20 h) [35] favored the transcyclization of the thiosuccinimide linker in the type 2 tryptic peptides.

Considering that proteolysis conditions also favored the hydrolysis of the thiosuccinimide linker [22, 33], we also search for type 2 tryptic peptides with the hydrolyzed thiosuccinimide linker (structures (**III**) and (**IV**), **Figure 1**) [20]. Peptides with this linker form also contain valuable information on the conjugation sites [20, 23]. Twenty-four MS/MS spectra were assigned to ten type 2 peptides with the hydrolyzed thiosuccinimide linker (**Table 3** and **Supplemental Files 7-8**). The MS/MS spectrum of the type 2 peptide ([^775^IIDYEY**<u>K</u>**IYSGPDK^768^]-(^538^CVNF^541^-H_6_)) shown in **Figure 3** is one example. The linker fragment ion C+98 (*m/z* = 701.784, 2+ and *m/z* = 1402.566, 1+) was generated by a favorable fragmentation of the newly formed pseudopeptide bond [23], which provided direct information on the molecular masses of the *C*-terminal peptide of the RBD (^538^**<u>C</u>**VNF^541^-HHHHHH) and the molecular mass of the crosslinked tryptic peptide of the d-TeNT (^775^IIDYEY**<u>K</u>**IYSGPDK^768^) can be inferred by the mass difference to the precursor ion. The thioether bond fragment ion named as P+203 (*m/z* = 636.287, 2+) [23] also confirmed the molecular mass of the d-TeNT crosslinked tryptic peptide ^775^IIDYEY**<u>K</u>**IYSGPDK^768^. In particular C+98 and P+203 ions are linker fragment ions exclusive for the hydrolyzed thiosuccinimide form. The identification of 10 conjugation sites based on type 2 peptides with the hydrolyzed thiosuccinimide linker (**Table 3**) did not represent a new contribution in addition to the 18 conjugation sites already assigned when considering the transcyclized linker (**Table 2**). However, twenty-four new MS/MS spectra (**Table 3**) provided additional support for the identification of 10 of the 18 conjugation sites shown in **Table 2**. Beside the quality of the MS/MS spectra, and the data validation supported by the linker fragment ions there are other experimental evidences that support the identifications of the type 2 peptides shown **Table 2** and **Table 3**:

1. Sixty-seven and seventy-five percent of all MS/MS spectra assigned to type 2 peptides with the transcyclized and hydrolyzed thiosuccinimide linkers, respectively, were coincidently assigned by both pLink2 [25] and Kojak [26] software (**Table 2**, **Table 3**, and **Supplemental file1, Fig. S15-S16**).
2. The charge state of all type 2 tryptic peptides regardless of linker type (hydrolyzed or transcyclized) was equal to or higher than 3+ (**Figure 4**). Eighty-one and eighty percent of all type 2 tryptic peptides with the transcyclized and hydrolyzed linker, respectively, were ionized with either 4+ or 5+. This behavior differs from the linear peptides which ionized preferably with 2+ and 3+ in 85 % of the cases. (**Figure 4**). This result is consistent with a previous report in the literature [51] despite of the *C*-terminal peptide of RBD is not a tryptic peptide with lysine or arginine residues at the *C*-terminus. The six histidine residues of the *C*-terminal peptide of RBD are responsible for efficiently retaining various protons in the gas phase in the same way that the *C*-terminal lysine or Arg does in tryptic peptides.
3. Beside the presence of specific linker fragment ions in the MS/MS spectra, we observed intense singly-charged ions at *m/z* = 110. 072, *m/z* = 275.126 and *m/z* = 412.185; which were not automatically assigned by the software. The first one corresponds to the immonium ion of histidine, and the others are internal fragment ions (HH and HHH) of the tandem repeat of six histidine residues present in all type 2 tryptic peptides identified here (**Table 2** and **Table 3**).

**Figure 3.**
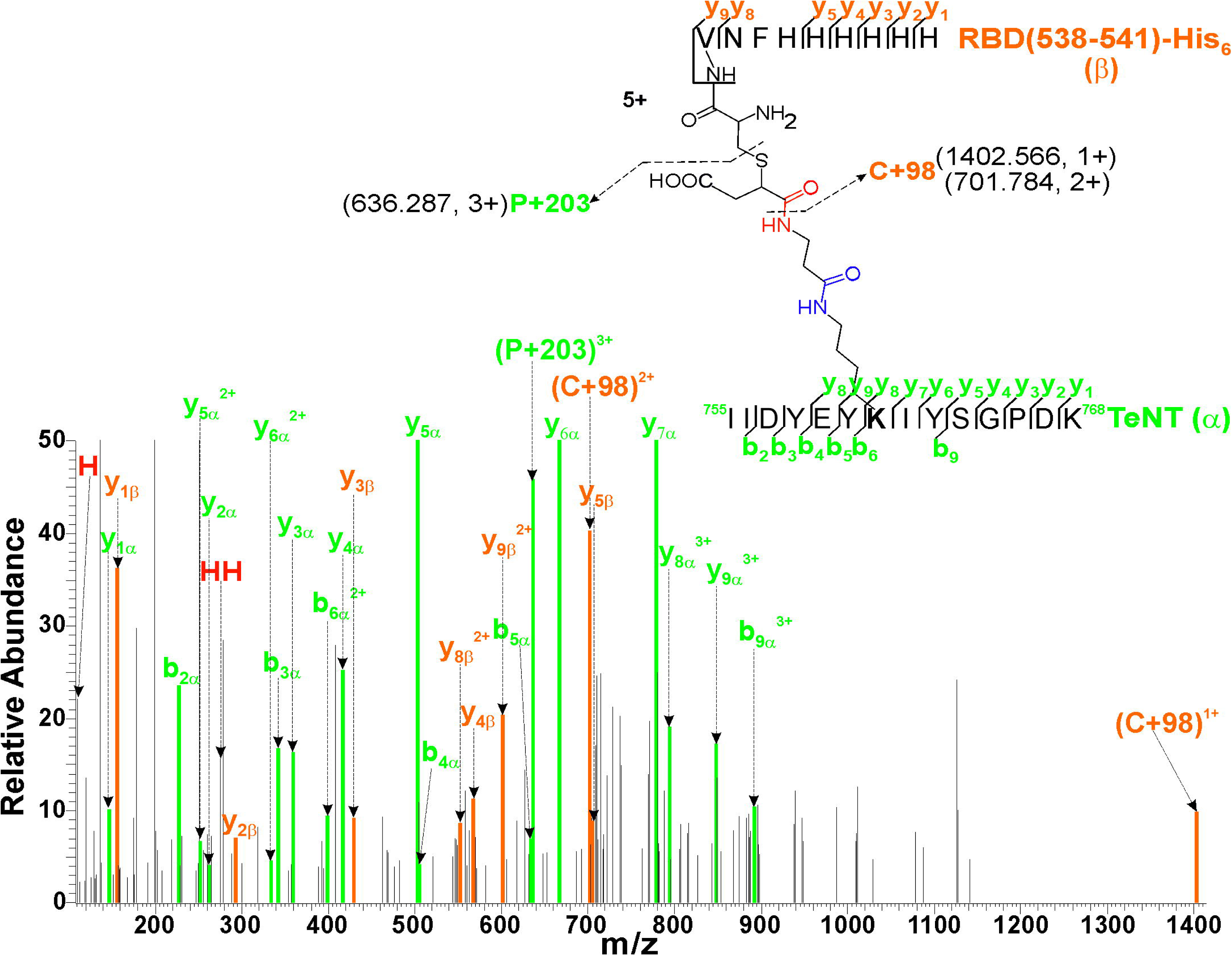
ESI-MS/MS spectrum assigned by pLink2 software [25] to a tryptic type 2 [^755^IIDYEY**<u>K</u>**IYSGPDK^768^]-(^538^**<u>C</u>**VNF^541^-HHHHHH) is composed by two peptides, one derived from the *C*-terminal peptide of the RBD of SARS-CoV-2 (^538^**<u>C</u>**VNF^541^-HHHHHH) and the other [^755^IIDYEY**<u>K</u>**IYSGPDK^768^] from the d-TeNT (UPI000003617E) of *C. tetani*. Amino acids highlighted in bold (C^538^ and K^761^) are crosslinked *via* a hydrolyzed linker. The (C+98) linker fragment ion detected at (*m/z* = 1402.566, 1+, *m/z* = 701.784, 2+), corresponds to the fragmentation of the pseudopeptide bond [22, 23] newly formed due to the hydrolysis reaction (highlighted in red in Figure 1, structures (**IV**) and (**V**)) [32]. The linker fragment ions detected at *m/z* = 636.287, 3+ corresponds the P+203, a fragment ion specific for the hydrolyzed thiosuccinimide linker. The signals at *m/z* = 110.071, 1+ and *m/z* = 275.125, 1+ correspond to the immonium ion of the histidine (H) and to an internal fragment ion (HH) of the tandem repeat of six histidine residues present in type 2 tryptic peptides identified here, respectively.

**Figure 4.**
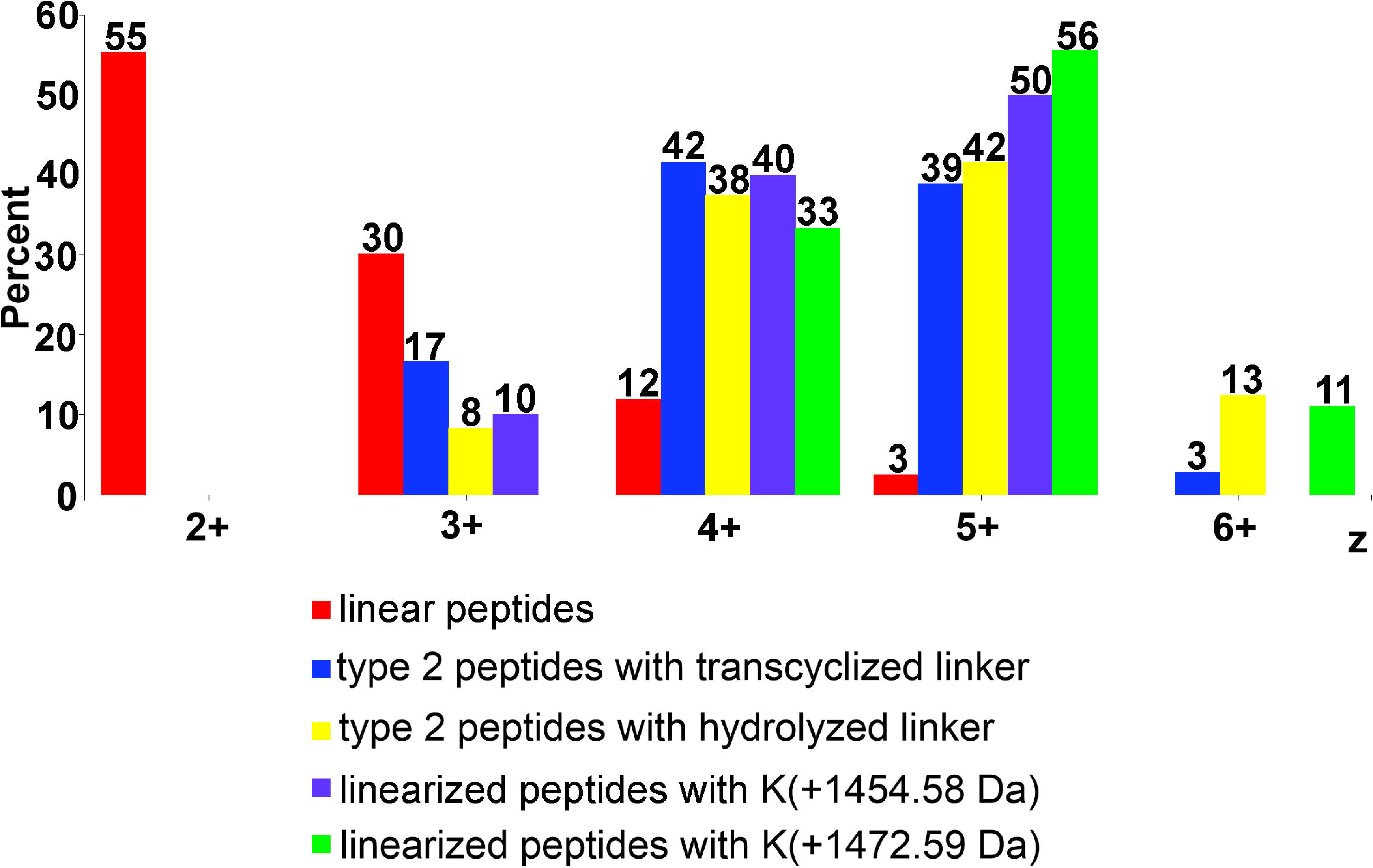
Charge state distribution expressed in percent for linear peptides, linearized peptides containing conjugated lysine residues (Δm= +1454.58 Da or Δm= +1472.59 Da) and type 2 tryptic peptides with the linker in the transcyclized and hydrolyzed forms. All peptides used to construct this bar graph were generated by tryptic digestion of d-TeNT and analyzed by LC-MS/MS.

**Table 3.**
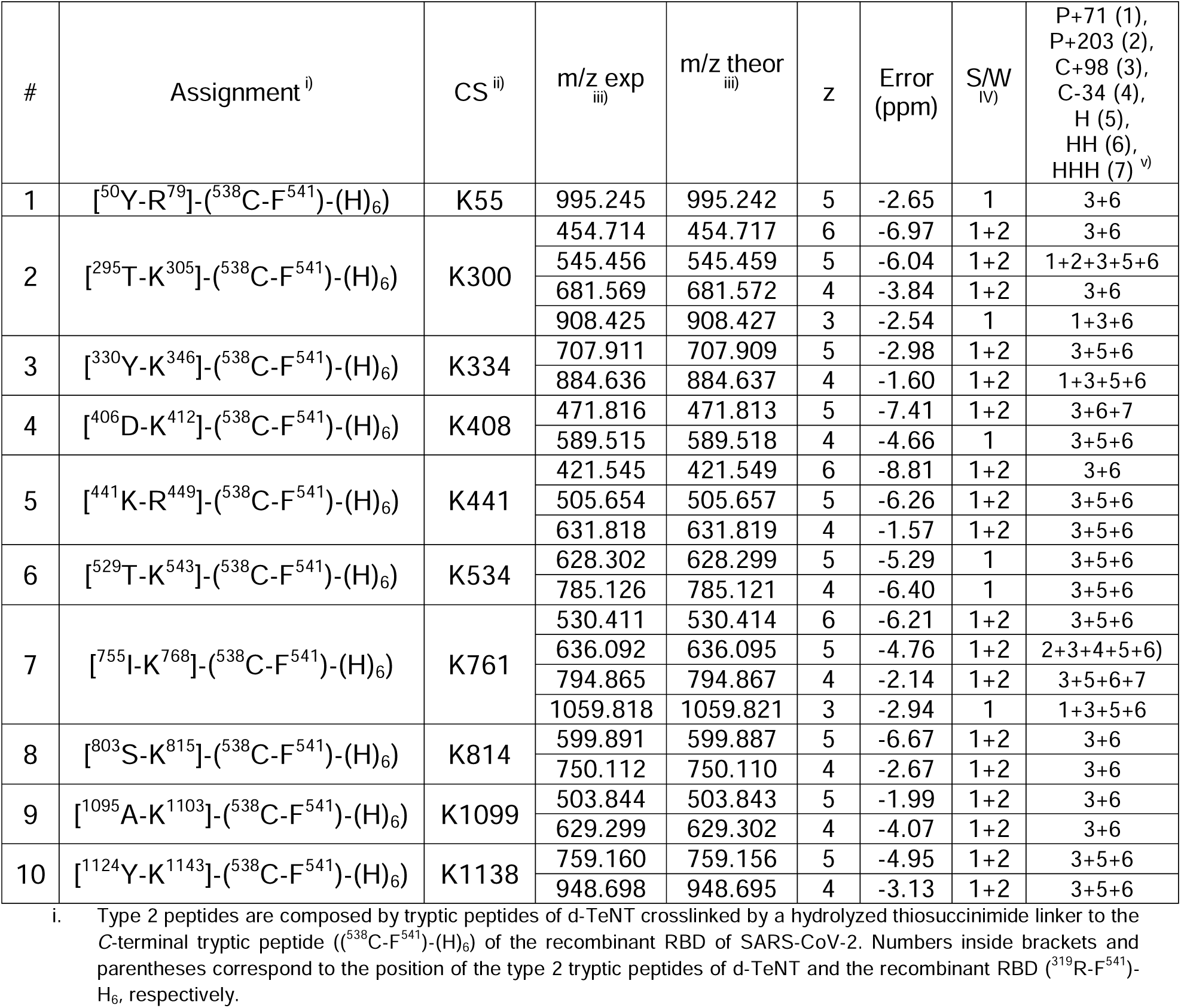
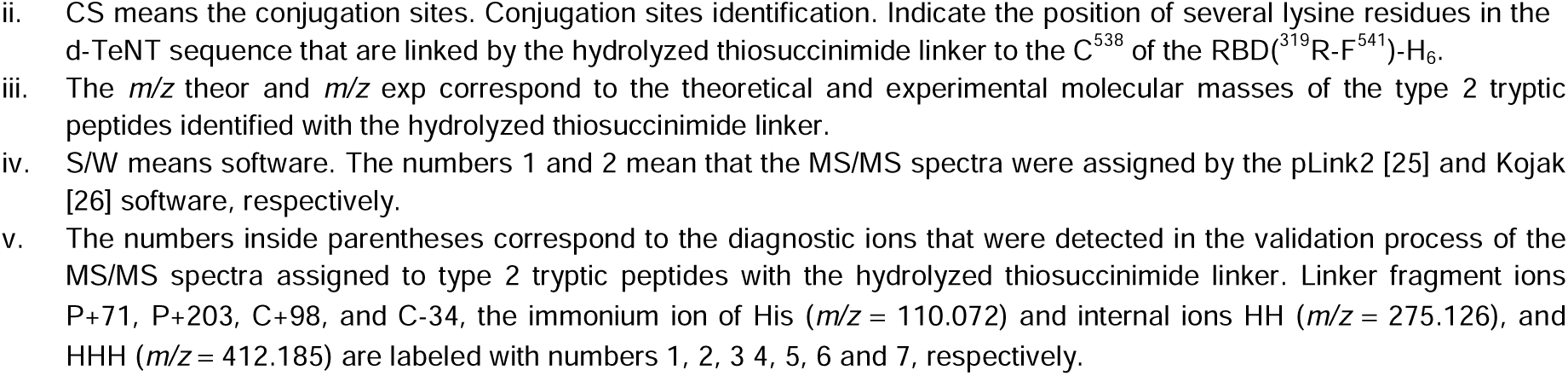
Conjugation sites identification in the d-TeNT (UPI000003617E) determined by LC-MS/MS analysis and based on the assignment of type 2 peptides with a hydrolyzed thiosuccinimide linker.

The validation approach described above makes the assignment of 60 MS/MS spectra to type 2 tryptic peptides and identification of the 18 conjugation sites a very high reliability (**Table 2** and **Table 3**).

We also based the identification of the conjugation sites of the d-TeNT following a linearization strategy. PEAKS Studio software [44] identified linear peptides carrying within their sequences conjugated lysine residues with a molecular mass increased by 1454.58 Da and 1472.59 Da when they were linked to the RBD *C*-terminal tryptic peptide through a transcyclized and hydrolyzed linker, respectively (**Table 4**). This strategy allowed the identification of twelve conjugation sites, being eight of them coincidently identified by Kojak and pLink2 software (**Fig. S17**). In the linearization strategy, four new conjugation sites were identified. Overlapping the information of provided by the identification of cross-linked peptides (**Table 2** and **Table 3**) and linearized cross-linked peptides (**Table 4**) a total of 22 conjugation sites for the d-TeNT were identified (**Fig. S17**).

**Table 4.**
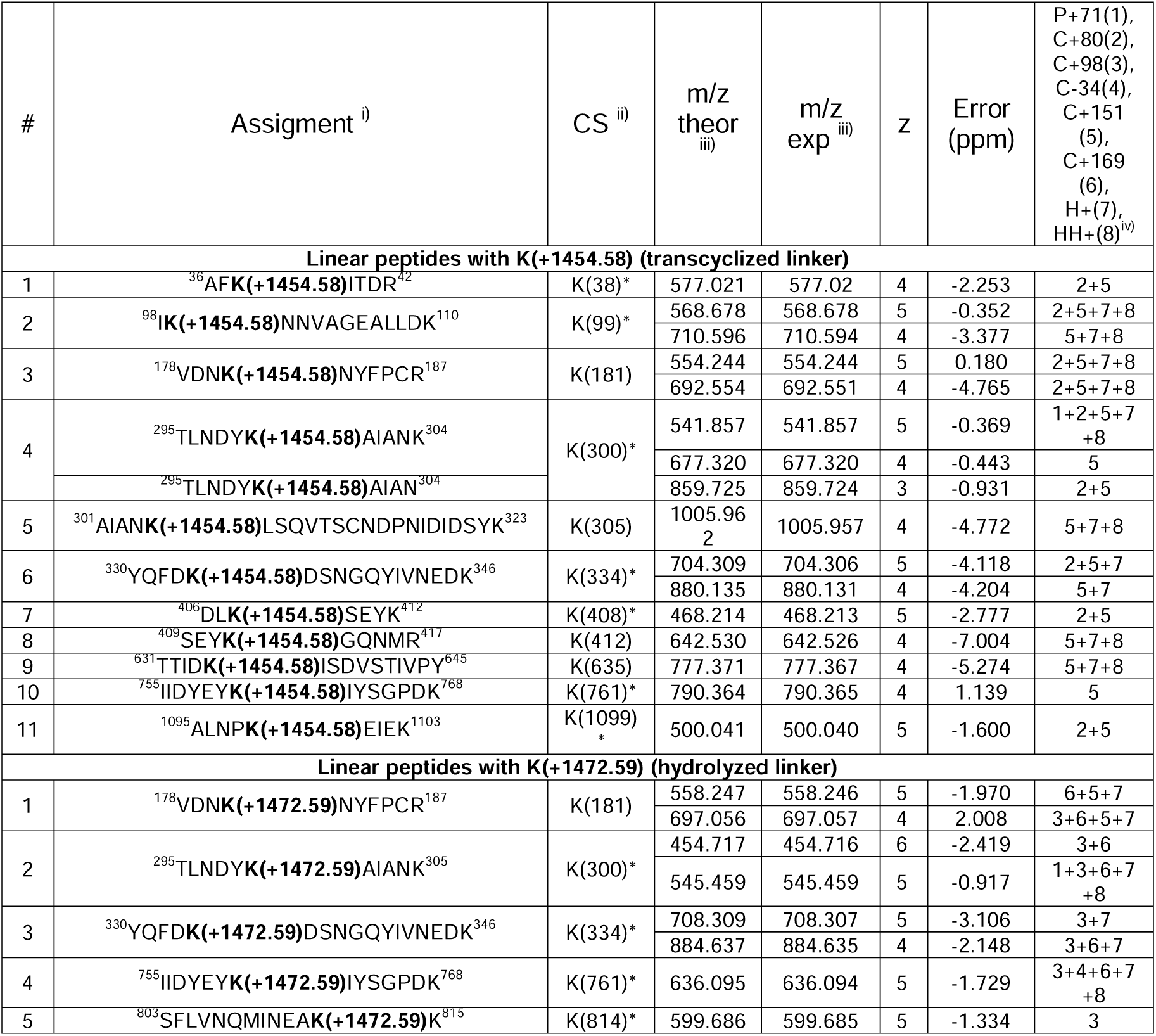
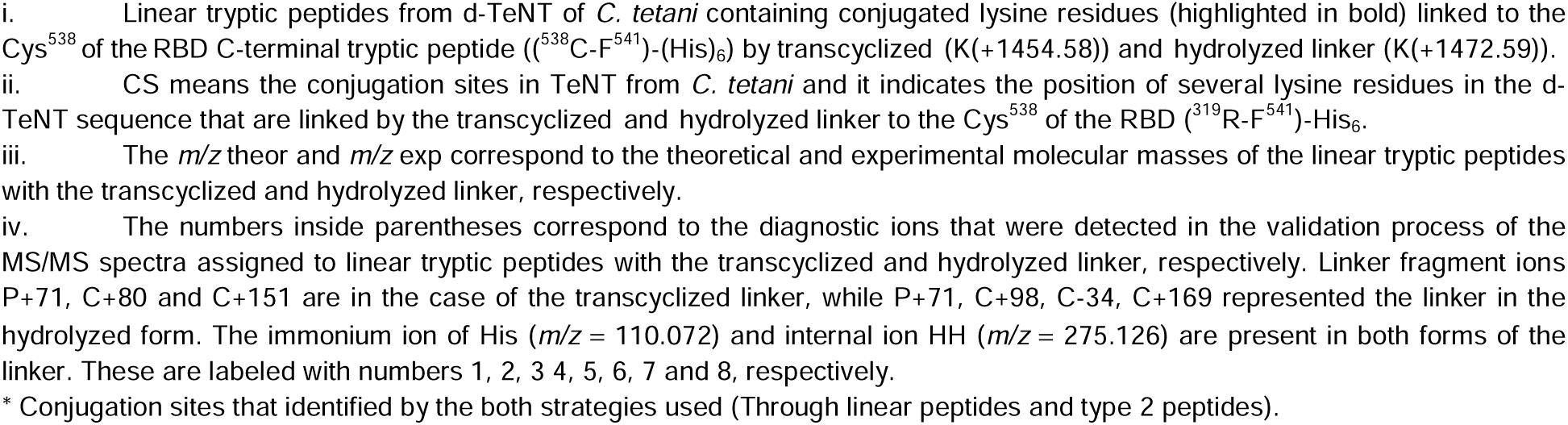
Conjugation sites in d-TeNT identified by PEAKS Studio [44] by MS/MS analysis. This assignment is based on linear peptides containing conjugated lysine residues linked to the RBD *C*-terminal tryptic peptide by a transcyclized (K(+1454.58)) and hydrolyzed (K(+1472.59)) linker forms.

Despite of any restriction on the charge state was considered during database search by using PEAKS [44], the charge state distribution of linearized peptides with (K+1454.58) and (K+1472.59) was also equal to or higher than 3+ (**Table 4**). This distribution resembles the one showed by cross-linked peptides (**Figure 4**) identified by pLink2 [25] and Kojak [26] software (**Table 2** and **Table 3**). It confirms that their structures, although identified as linear peptides with (K+1454.58) and (K+1472.59) by PEAKS Studio [44], they are cross-linked peptides.

The identification of the conjugation sites based on the assignment of the MS/MS spectra to type 2 peptides was a strategy more efficient than the one based on the linear peptides carrying both linker forms (**Supplemental file 1**, **Fig. S17** and **Fig. S18**). In addition, the transcyclization reaction seems to be more favored than the hydrolysis of the linker. Twenty-two and eleven conjugation sites were identified by crosslinked peptides carrying a transcyclized and hydrolyzed linker, respectively (**Supplemental file 1**, **Fig. S19**). All conjugation sites identified by peptides with the hydrolyzed linker were also identified by peptides with the transcyclized linker (**Supplemental file 1**, **Fig. S19**). The MS/MS spectra assigned to linearized peptides were validated following the same criteria used for the identification of cross-linked peptides. These results support that the strategy for identifying linearized peptides by PEAKS was also successful for the identification of the conjugation sites in the d-TeNT.

The pLink2 [25] and Kojak [26] software also assigned other forty-five and twenty-four MS/MS spectra to type 2 tryptic peptides derived from ten other proteins different to d-TeNT with the transcyclized (**Supplemental Table S1**) and hydrolyzed (**Supplemental Table S2**) linkers, respectively. The MS/MS spectra of two type 2 peptides shown in **Fig. S20-S21 (Supplemental file 1),** confirm this assessment. For UPI0002DAC767, the second most abundant protein (see **Table 1**), nine conjugation sites were identified based on the assignment of MS/MS spectra to type 2 peptides with the two linkers forms (**Supplemental Table S1** and **S2**). For proteins, UPI00001A9618, UPI000001030F and UPI000001056B only one conjugation site was identified, although they have several lysine within their sequences (**Table 1** and **Supplemental Tables S1** and **S2**). The abundance of these proteins is less than 5% for the TT preparation analyzed (**Table 1**) and probably it affects the detection of their conjugation sites, despite the enrichment step of crosslinked peptides prior to LC-MS/MS analysis. The identification of conjugation sites in other proteins different to d-TeNT were also supported by linearized peptides carrying conjugated lysine residues ((K+1454.58) and (K+1472.59)) as shown in **Supplemental Table S3**. The charge state distribution of peptides carrying conjugation sites of proteins other than d-TeNT, regardless of the strategy used for their identification, as linearized or cross-linked peptides, was similar to that of the type 2 peptides derived from d-TeNT (**Figure 4**).

## 4 Conclusions

The TT preparation used as a carrier in SOBERANA^®^02 is composed of a complex mixture of 909 proteins, of which d-TeNT is the most abundant one regardless of the quantification method used. The abundance of most of the other proteins in the TT preparation is less than 5%.

The activation process incorporated a maleimide group in 102 out of 107 of the lysine residues, which prove the high efficiency of this process. This result is important because it increases the chances for conjugating the new protein – and also putative peptide and glycan antigens – to T cell epitopes, a process that is often relevant for a more potent immune response.

Interestingly, twenty-two lysine residues in TT were found to be linked to the C^538^ in the RBD *C*-terminal peptide, suggesting that these residues are those better exposed in the protein surface to receive the addition of the bulky, 30 kDa RBD protein bearing a free thiol group. We also found the same RBD *C*-terminal peptide linked to lysine residues of other ten low-abundance abundant proteins present in the TT preparation.

The assignment of the conjugation sites was carried out with great reliability because all MS/MS spectra were validated following orthogonal criteria, such as the presence in the MS/MS spectra of specific linker fragment ions and diagnostic ions.

Whereas SOBERANA^®^02 seems to be the first clinically validated protein-protein conjugate vaccine, our results are important in the context of any conjugate vaccine based on TT as a carrier protein(s) mixture. Our successful characterization of this protein conjugate(s) by LC-MS/MS includes not only the reliable determination of the conjugation sites but also the minor protein components present in the TT preparations. Consequently, we truly believe this report is very relevant for the conjugate vaccine community since TT is considered worldwide as a highly effective carrier protein, which is currently used in various polysaccharide-protein conjugates but can also be useful in next-generation protein-protein conjugates.

## Supporting information

Supplemental file 2. Protein composition

Supplemental file 3. Maleimide activation process

Supplemental file 4. Activation process Maleimide plus cysteamin

Supplemental file 5. Type 2 peptides with transcyclized linker identified by pLink2

Supplemental file 6. Type 2 peptides with transcyclized linker identified by kojak

Supplemental file 7. Type 2 peptides with hydrolyzed linker identified by pLink2

Supplemental File 8. Type 2 peptides with hydrolyzed linker identified by kojak

## Acknowledgements

The authors thank FIOCRUZ for using their Network of Technological Platforms. This work was partially supported by the German Ministry of Education and Science (01DN18015) and the Max Planck Society for the Advancement of Science.

## CRediT

**OM:** Investigation, Validation, Data Curation, Formal Analysis, Visualization. **DS-M:** Investigation, Conceptualization. **SP:** Investigation. **JPS:** Investigation.**AR-U:** Formal Analysis. **PER-B:** Software. **VB:** Investigation, Funding Acquisition. **RG:** Investigation. **PC:** Software, Investigation, Funding acquisition. **MB**: Investigation. **KZ:** Investigation. **JW:** Investigation, Funding Acquisition. **TB**: Investigation, **YVB** Resources, Conceptualization, Project Administration, Supervision. **DG-R:** Writing - Review & Editing, Conceptualization, Investigation. **DGR:** Conceptualization, Writing - Review & Editing. **VVB**: Resources, Conceptualization, Project Administration, Supervision. **LJGL**: Investigation, Methodology, Supervision, Conceptualization, Writing - Original Draft, Writing - Review & Editing.

