## Supplemental file 1 for "LC-MS/MS characterization of SOBERANA^®^02, a receptor binding domain-tetanus toxoid conjugate vaccine against SARS-CoV-2": 01_Supplemental File 1.docx

**Title:** LC-MS/MS characterization of SOBERANA^®^02, a receptor binding domain-tetanus toxoid conjugate vaccine against SARS-CoV-2.

**Authors:** Olivia Martínez^1^, Darielys Santana-Medero^1^, Satomy Pousa^2^, Arielis Rodríguez-Ulloa^2^, Jean Pierre Soubal^1^, Pablo E. Ramos-Bermúdez^2^, Raine Garrido^1^, Vladimir Besada^2^, Paulo Carvalho^3^, Michel Batista^3^, Katharina Zetl^4^, Jacek Wiśniewski^4^, Tamy Boggiano^5^, Yury Valdés-Balbín^1^, Dagmar García-Rivera^1^, Daniel G. Rivera^1, 6^, Vicente Verez-Bencomo^1^, Luis Javier González ^2, ¥^.

**Affiliations:**

(1) Finlay Institute of Vaccines, 200 and 21 Street, Havana 11600, Cuba.

(2) Center for Genetic Engineering and Biotechnology, Ave 31 e/158 y 190, Havana 10600, Cuba.

(3) Carlos Chagas Institute/FioCruz Rua Professor Algacyr Munhoz Mader, 3775 CIC 81350-010, Curitiba, Brazil.

(4) Department Proteomics and Signal Transduction, Max Planck Institute of Biochemistry. Am Klopferspitz 18, 82152, Martinsried, Germany.

(5) Center for Molecular Immunology, P.O. Box 16040, 216 St., Havana, Cuba

(6) Laboratory of Synthetic and Biomolecular Chemistry, Faculty of Chemistry, University of Havana, Zapata and G, 10400, La Habana, Cuba.

**^(¥)^ Corresponding author:**

Luis Javier González López PhD

Mass Spectrometry Laboratory, Department of Proteomics.

Center for Genetic Engineering and Biotechnology.

Avenida 31, e/ 158 y 190, Cubanacán, Playa. CP 11600, PO. Box 6162, La Habana 11600, Cuba.

**Index**

**Fig. S1**. Activation process of the d-TeNT. Firstly, d-TeNT reacts with the heterobifunctional crosslinking reagent BMPS (3-maleimidopropionic acid *N*-hydroxysuccinimide) and several *N*-propionyl maleimide groups are added to primary amino groups of d-TeNT (at the Lys residues and the *N*-terminus) by increasing the molecular mass of the lysine residues by +151.03 Da (K(+151.03)). Eventually, the newly incorporated maleimide group during proteolysis can be hydrolyzed to yield another modified lysine residue with the molecular mass increased by 169.04 Da (K(+169.04)). In a second step, the maleimide group in the d-TeNT can react with cysteamine molecule by the Michael addition forming a thioether bond and a modified Lysine residue with its molecular mass increased by 228.06 Da (K(+228.06)). Eventually, this structure can be also hydrolyzed during the proteolysis to yield another modified lysine residue with the molecular mass increased by 246.07 Da (K(+246.07)).

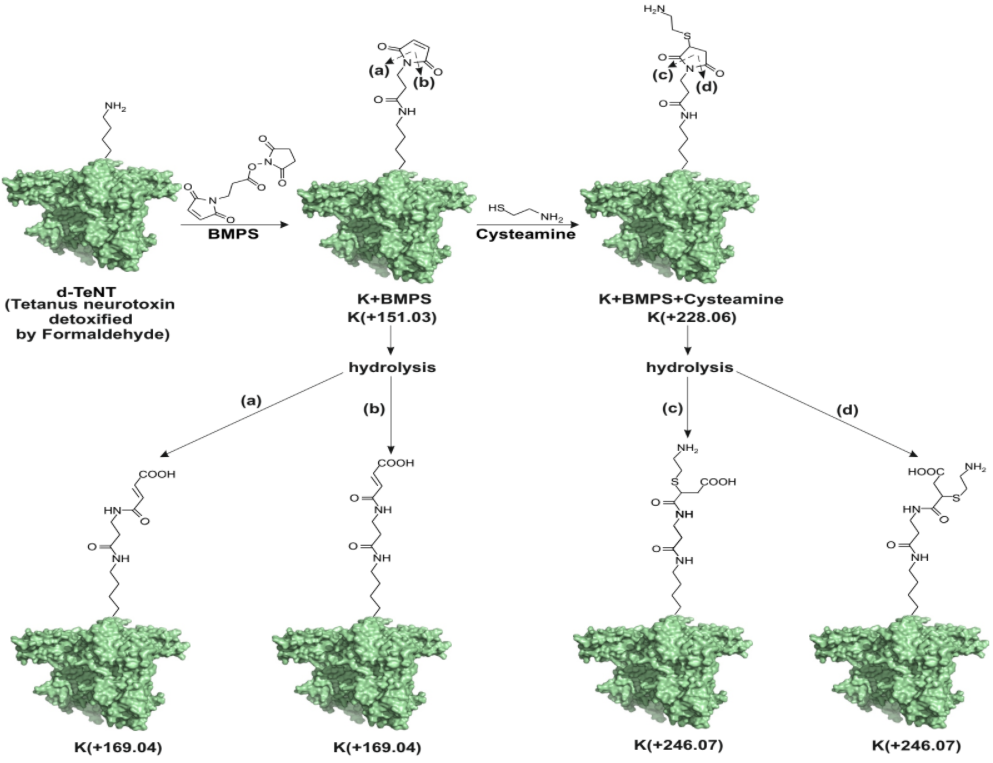

**Fig. S2**. Fragmentation of the newly formed pseudopeptide bond (highlighted in red) in a transcyclized linker between a lysine and cysteine residue. The Lysine and Cysteine residues in this structure represent two hypothetical peptides. The fragmentation of this pseudopeptide bond yields two linker fragment ions named here as P+71 and C+80 [1, 2].

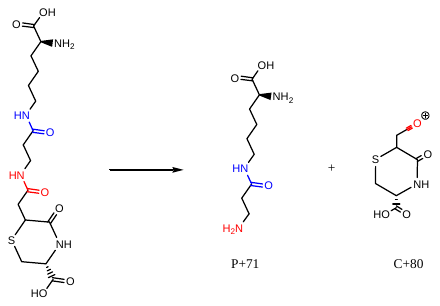

**Fig. S3**. Fragmentation of the newly formed pseudopeptide bond (highlighted in red) in a hydrolyzed thiosuccinimide linker between a lysine and cysteine residue. The Lysine and Cysteine residues in this structure represent two hypothetical peptides. The fragmentation of this pseudopeptide bond is a metastable fragmentation that yields two linker fragment ions named here as P+71 and C+98 [1].

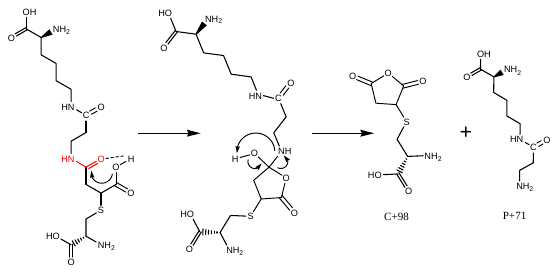

**Fig. S4.** Fragmentation of the pseudopeptide bond present in the thiosuccinimide (structure **III**) and all its stabilized forms: hydrolyzed thiosuccinimide linker (structures **I** and **II**) and thiazine (structure **IV**). All linker forms upon fragmentation in gas phase could yield the “P ions” corresponding to the unmodified peptide with the crosslinked lysine when protonated (structure **V**). In addition, the isomeric species containing the hydrolyzed thiosuccinimide linker yield the C+169 ions corresponding to the peptide with the crosslinked Cysteine residue involved in the thioether bond (structure **VI** and **VII**). The two isobaric linker forms, thiosuccinimide and transcyclized linker, yield the C+151 ions but with two different structures as shown in the figure (structures **VIII** and **IX**).

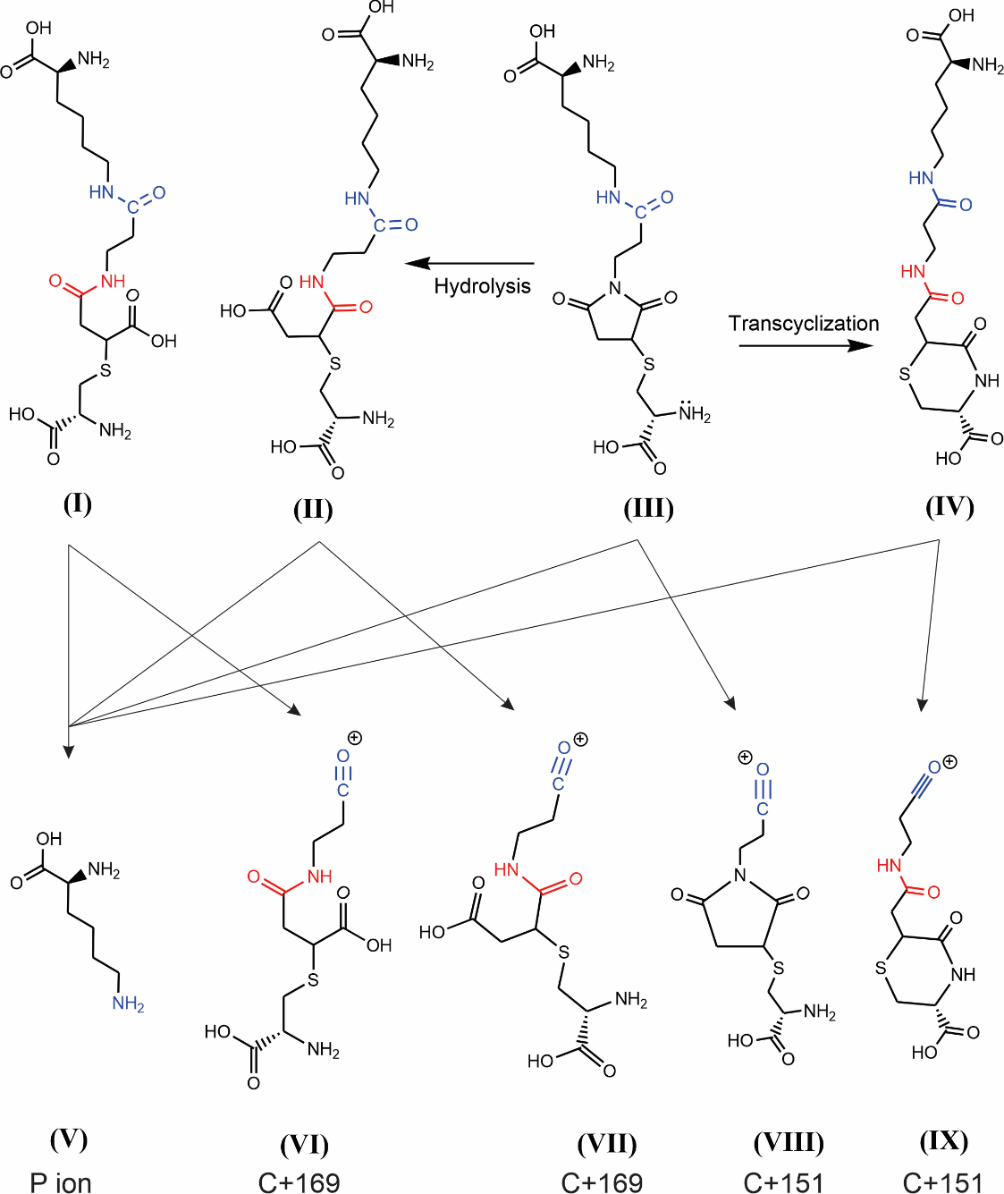

**Fig. S5**. Fragmentation of the thioether bond in a hydrolyzed thiosuccinimide linker between a lysine and cysteine residue. The lysine and cysteine residues in this structure represent two hypothetical peptides, P1 and P2, crosslinked by one of the two possible positional isomers of the hydrolyzed thiosuccinimide linker ((P1-P2)_H_). The fragmentation of the thioether bond by a beta elimination and a proton transfer yields two linker fragment ions designed here as P1+203 Da and the P2-34 Da.

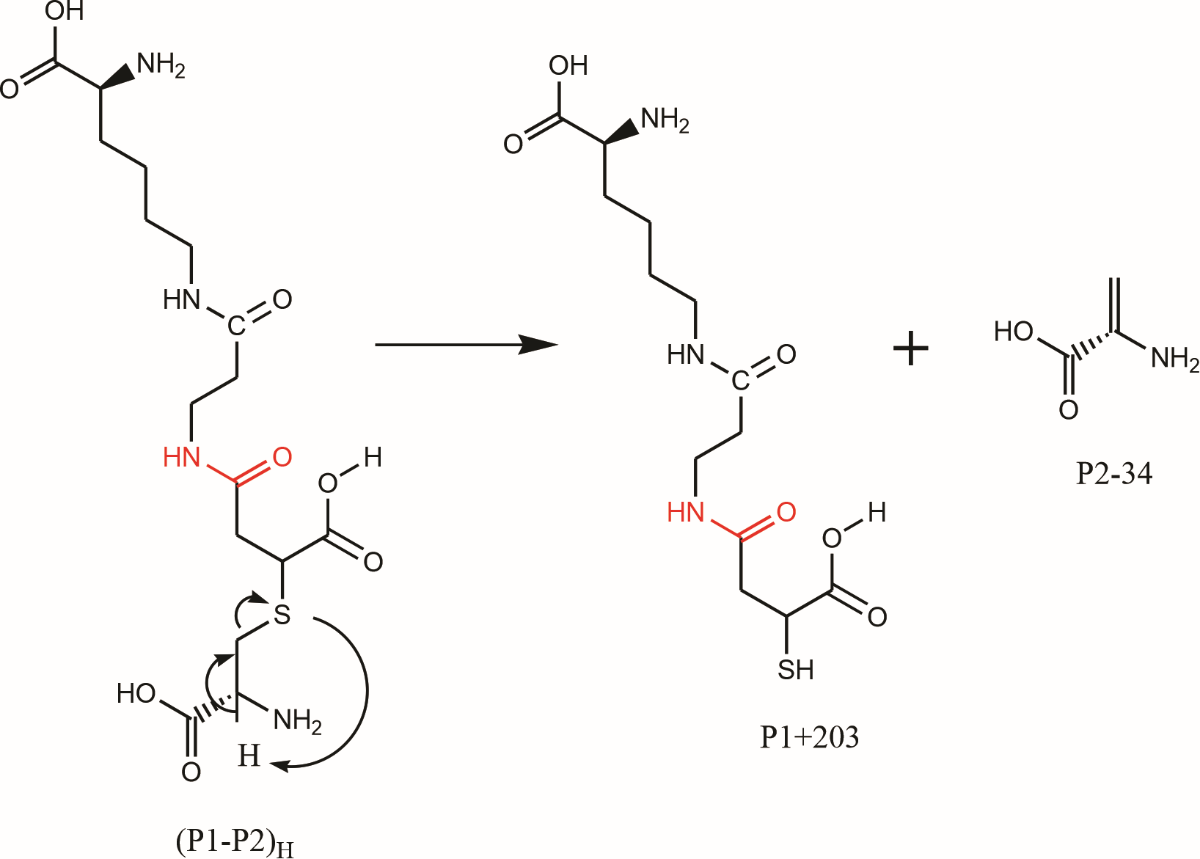

**Fig. S6.** Fragmentation of the thioether bond in a hydrolyzed thiosuccinimide linker between a lysine and cysteine residue. The Lysine and Cysteine residues in this structure represent the P1 and P2 peptides crosslinked by one of the positional isomers of the hydrolyzed thiosuccinimide linker ((P1-P2)_H_). The fragmentation of the thioether bond by a beta elimination and a proton transfer yields two linker fragment ions designed here as P1+169 Da and the reduced P2-SH.

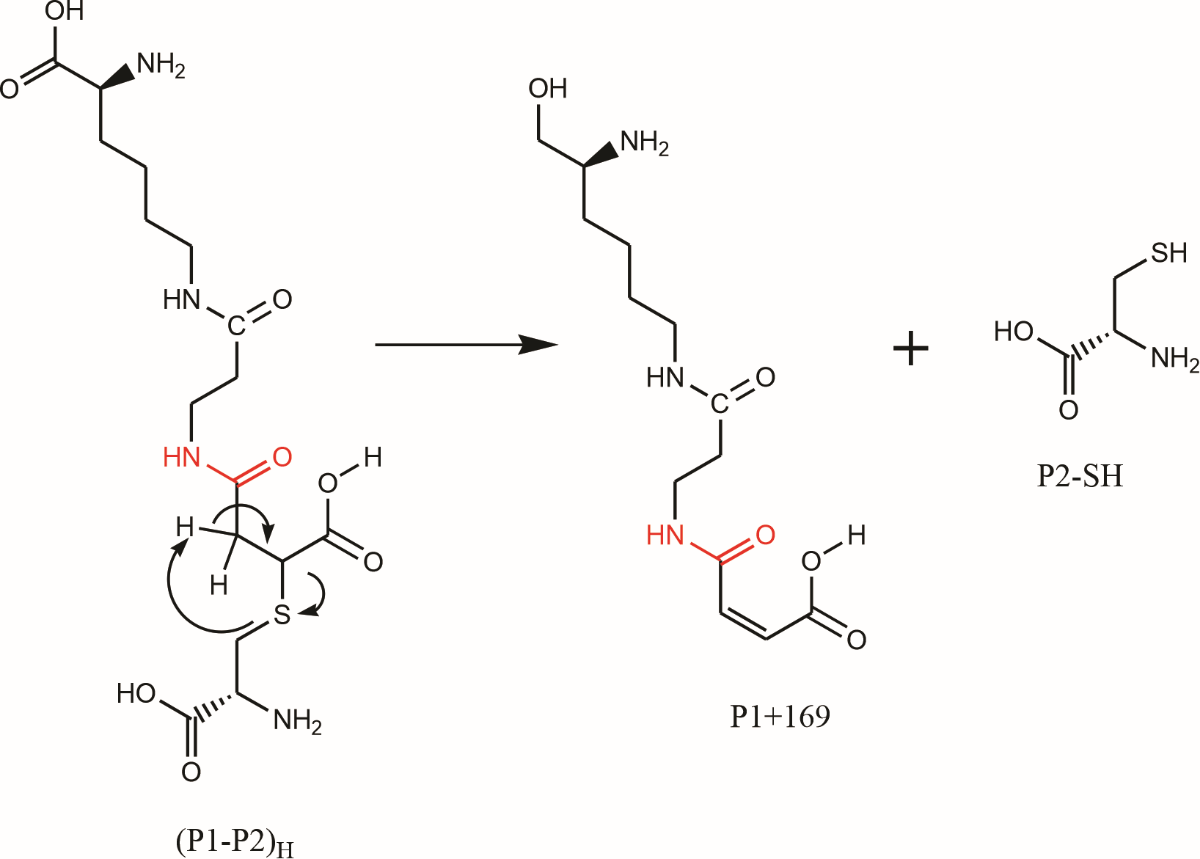

**Fig. S7.** Proportional Venn Diagram [3] overlaps the total proteins identified by PEAKS Studio V7.0 [4] and MaxQuant (v2.0.3.1) [5] software when tetanus toxoid preparation used in the synthesis of SOBERANA^®^02 was digested with Lys-C, trypsin and chymotrypsin using the MED-FASP protocol [6] and analyzed by LC-MS/MS. *C. tetani* protein database (UP000290279) containing 2708 entries (downloaded 13^th^ June 2024, https://www.uniprot.org) was queried and 1% FDR was accepted in the protein identification output.

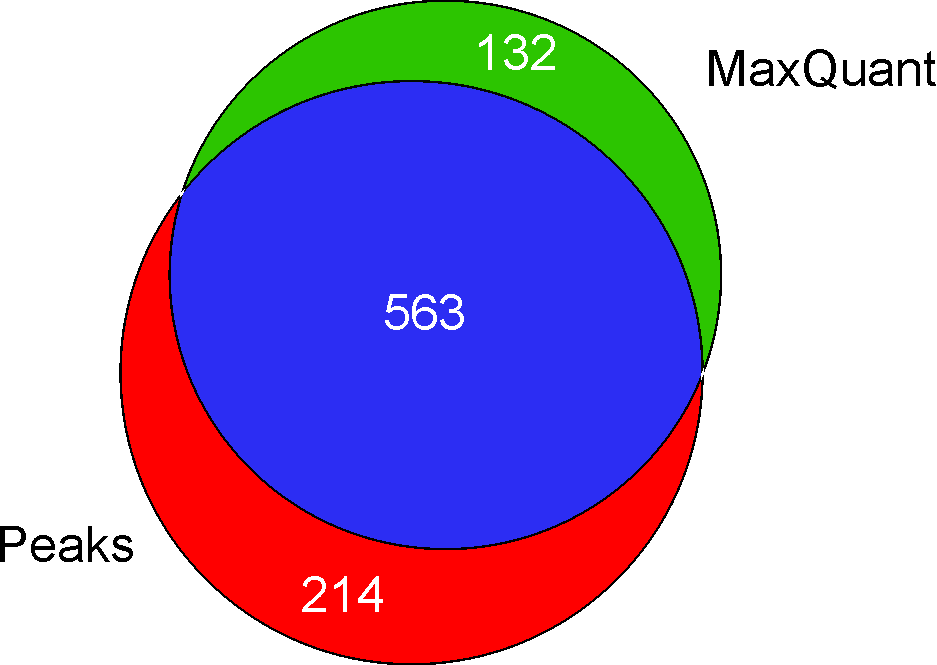

**Fig. S8**. Proportional Venn Diagram [3] overlapping the proteins identified by MaxQuant (v2.0.3.1) [5] software when tetanus toxoid preparation used in the synthesis of SOBERANA^®^02 was digested with Lys-C, trypsin and chymotrypsin (CT) using the MED-FASP protocol [6] and analyzed by LC-MS/MS. *C. tetani* protein database (UP000290279) containing 2708 entries (downloaded 13^th^ June 2024, https://www.uniprot.org) was queried and 1% FDR was accepted for the protein identifications.

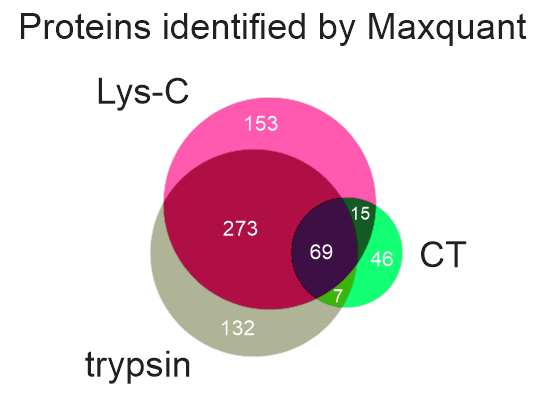

**Fig. S9**. Proportional Venn Diagram [3] overlapping the proteins identified by PEAKS Studio V7.0 [4] software when tetanus toxoid preparation used in the synthesis of SOBERANA^®^02 was digested with Lys-C, trypsin and chymotrypsin (CT) using the MED-FASP protocol [6] and analyzed by LC-MS/MS. *C. tetani* protein database (UP000290279) containing 2708 entries (downloaded 13^th^ June 2024, https://www.uniprot.org) was queried and 1% FDR was accepted for the protein identifications.

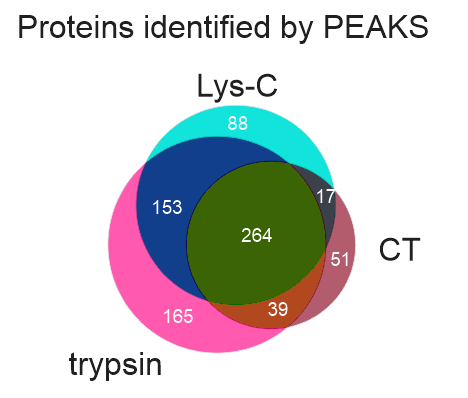

**Fig. S10**. Proportional Venn Diagram [3] showing the lysine residues found in the LC-MS/MS analysis of the detoxified tetanus neurotoxin ([UPI000003617E](https://www.uniprot.org/uniparc/UPI000003617E/entry), d-TeNT) acylated with either the intact (K(+151.03)), or hydrolyzed (K(+169.04)) *N*-propionylmaleimide group after considering the information provided by considering all the information provided by Lys-C, trypsin and chymotrypsin proteolytic digestions [6]. The figures in the Venn diagram show the number of lysine residues modified with *N*-propionyl maleimide groups (intact and hydrolyzed) in the most-abundant protein ([UPI000003617E](https://www.uniprot.org/uniparc/UPI000003617E/entry)).

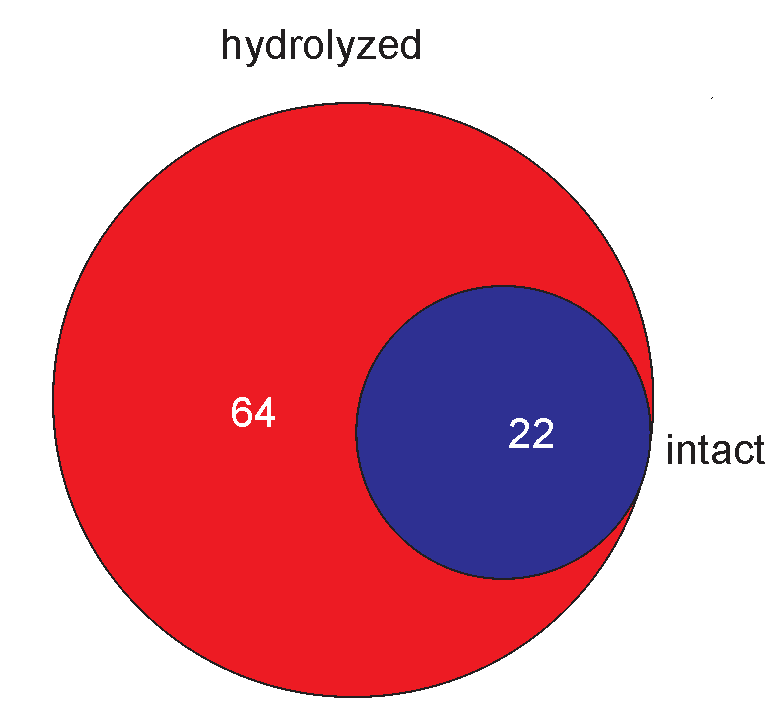

**Fig. S11**. The proportional Venn Diagram [3] shows the number of lysine residues that were found in the LC-MS/MS analysis of the detoxified tetanus neurotoxin ([UPI000003617E](https://www.uniprot.org/uniparc/UPI000003617E/entry), d-TeNT) modified with either the intact or hydrolyzed *N*-propionyl maleimide residue. The information provided here summarizes the individual contribution of three proteolytic digestions (Lys-C, trypsin and chymotrypsin (CT)) [6].

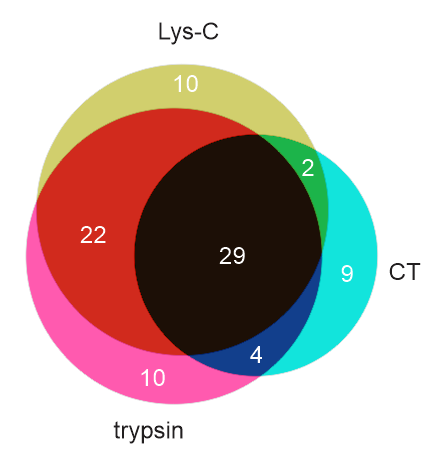

**Fig. S12.** A proportional Venn Diagram [3] overlapping the total number of lysine residues found in the LC-MS/MS analysis of the detoxified tetanus neurotoxin ([UPI000003617E](https://www.uniprot.org/uniparc/UPI000003617E/entry), d-TeNT) that were modified by incorporating N-propionylmaleimide group and a cysteamine molecule (NH_2_-CH_2_-CH_2_-SH) and during the MED-FASP digestion protocol [6] were detected in the intact ((K+228.06)) and the hydrolyzed form ((K+246.07)).

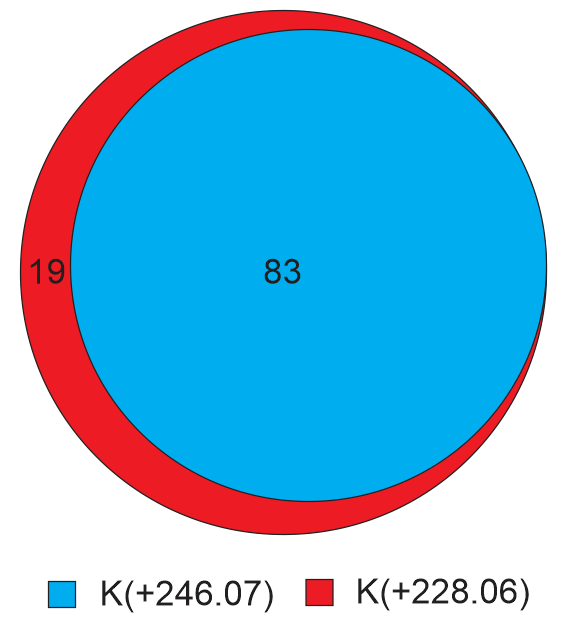

**Fig.S13**. A proportional Venn Diagram [3] overlapping the total number of lysine residues found in the LC-MS/MS analysis of the Lys-C, trypsin and chymotrypsin digestions [6] of the detoxified tetanus neurotoxin ([UPI000003617E](https://www.uniprot.org/uniparc/UPI000003617E/entry), d-TeNT) that were modified by incorporating *N*-propionylmaleimide group and a cysteamine molecule (NH_2_-CH_2_-CH_2_-SH) that were detected with the intact thiosuccinimide linker ((K+228.06)).

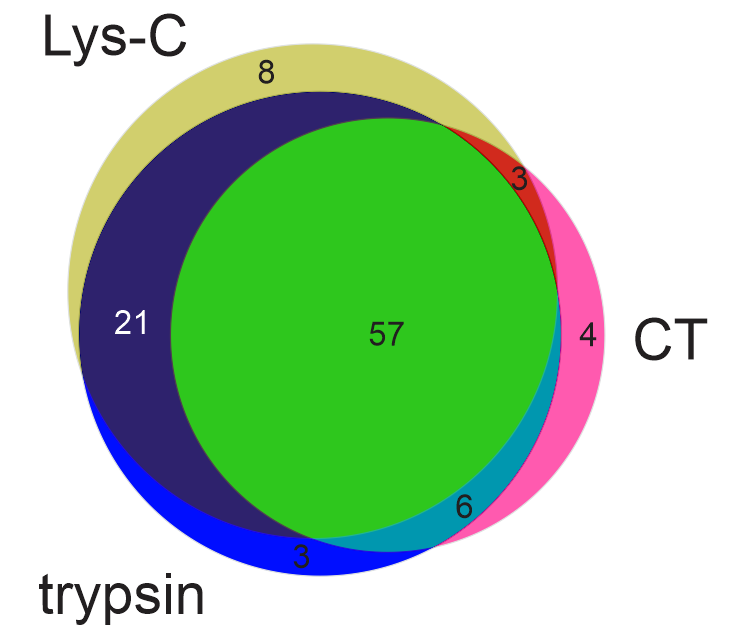

**Fig. S14**. A proportional Venn Diagram [3] overlapping the total number of lysine residues found in the LC-MS/MS analysis of the Lys-C, trypsin and chymotrypsin digestions [6] of the detoxified tetanus neurotoxin ([UPI000003617E](https://www.uniprot.org/uniparc/UPI000003617E/entry), d-TeNT) that were modified by incorporating *N*-propionylmaleimide group and a cysteamine molecule (NH_2_-CH_2_-CH_2_-SH) that were detected with the hydrolyzed thiosuccinimide linker ((K+246.07)).

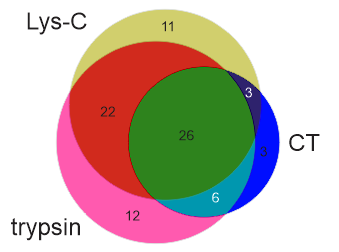

**Fig. S15.** Proportional Venn diagram [3] showing the overlapping results obtained by pLink2 [7] and Kojak [8] software for assigning the type 2 peptides with the transcyclized linker.

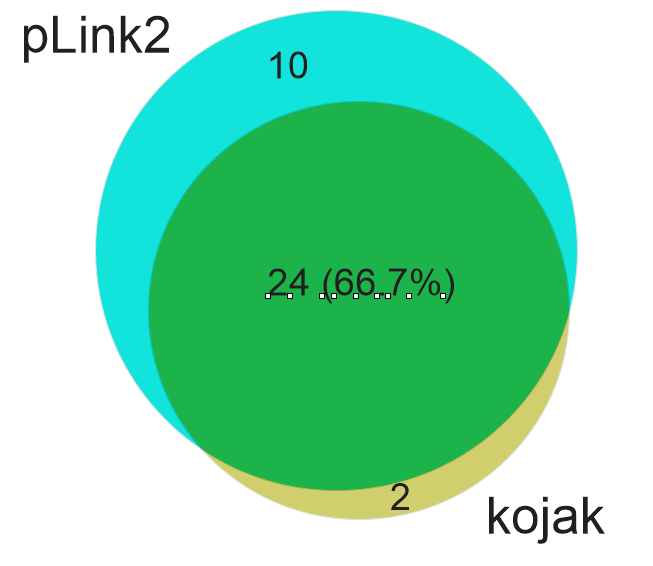

**Fig. S16.** Proportional Venn diagram [3] showing the overlapping results obtained by pLink2 [7] and Kojak [8] software for assigning the type 2 peptides with the hydrolyzed linker.

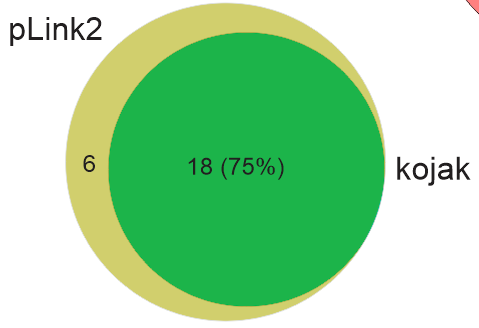

**Fig. S17**. Proportional Venn diagram showing the assignment of twenty-two conjugation sites determined by LC-MS/MS analysis of SOBERANA^®^02 tryptic peptides enriched after Ni^2+^-NTA chromatography. The identification of the conjugation sites was based on the assignment of the MS/MS spectra to linear and type 2 peptides, independent of the type of linker, either in hydrolyzed or transcyclized forms.

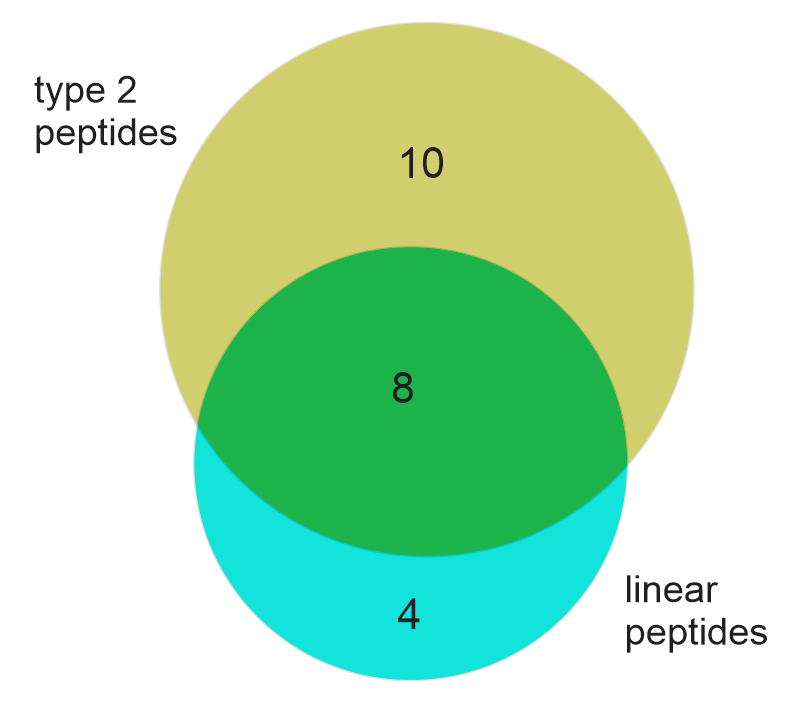

**Fig. S18.** Proportional Venn diagrams showing the identification of twenty-two conjugation sites in SOBERANA^®^02 as determined by LC-MS/MS analysis. Two strategies were used to obtain this result. One strategy was based on the assignment of MS/MS spectra to type 2 peptides and linear peptides with the transcyclized linker (see Venn diagram on the left), while the other strategy considered the contribution of the same peptides with the hydrolyzed linker (see Venn diagram on the right). For the identification of linear peptides containing conjugation sites with the transcyclized and hydrolyzed linkers, the molecular masses of the lysine residues were increased by (+1454.58 Da) and (+1472.59 Da), respectively.

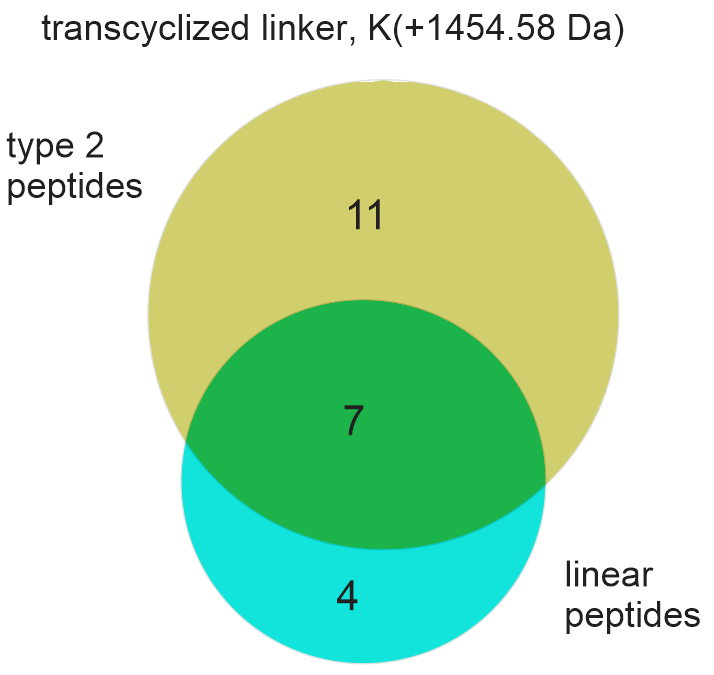

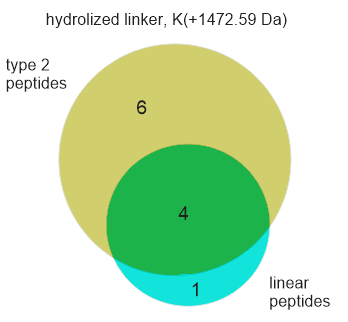

**Fig. S19**. Proportional Venn diagram showing the overlapping results for the assignment of the conjugation sites determined by the assignment of the MS/MS spectra to linear and type 2 peptides with the transcyclized and hydrolyzed linkers in SOBERANA^®^02.

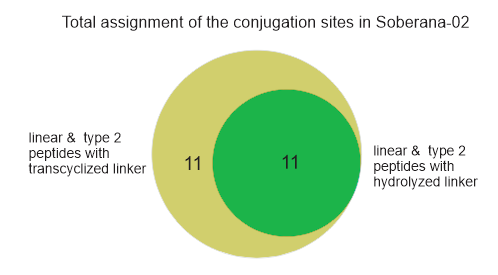

**Fig S20.** ESI-MS/MS spectrum assigned by pLink2 software [7] to a tryptic type 2 peptide composed by the *C*-terminal peptide of the RBD of SARS-CoV-2 (^538^**C**VNF^541^-HHHHHH) cross-linked via a transcyclized linker to the peptide [^338^**K**EIEDR^343^] of Chaperonin GroEL, Chaperonin-60, Cpn60 (UPI00016089BF) from *C. tetani*. This cross-linked peptide was detected at *m/z*= 561.754, 4+. The (P+71) and (C+80) linker fragment ions detected at (*m/z* = 430.727, 2+) and (*m/z* = 692.777, 2+ and *m/z* = 462.189, 3+), respectively were generated by the fragmentation of the pseudopeptide bond (highlighted in red) [9] newly formed due to the transcyclization reaction [10, 11]. The signals at *m/z* = 110.064 (1+) and *m/z* = 275.126 (1+) corresponds to the immonium ion of the histidine (H) and an internal fragment ion (HH) of the tandem repeat of six histidine residues present in type 2 tryptic peptides identified here, respectively.

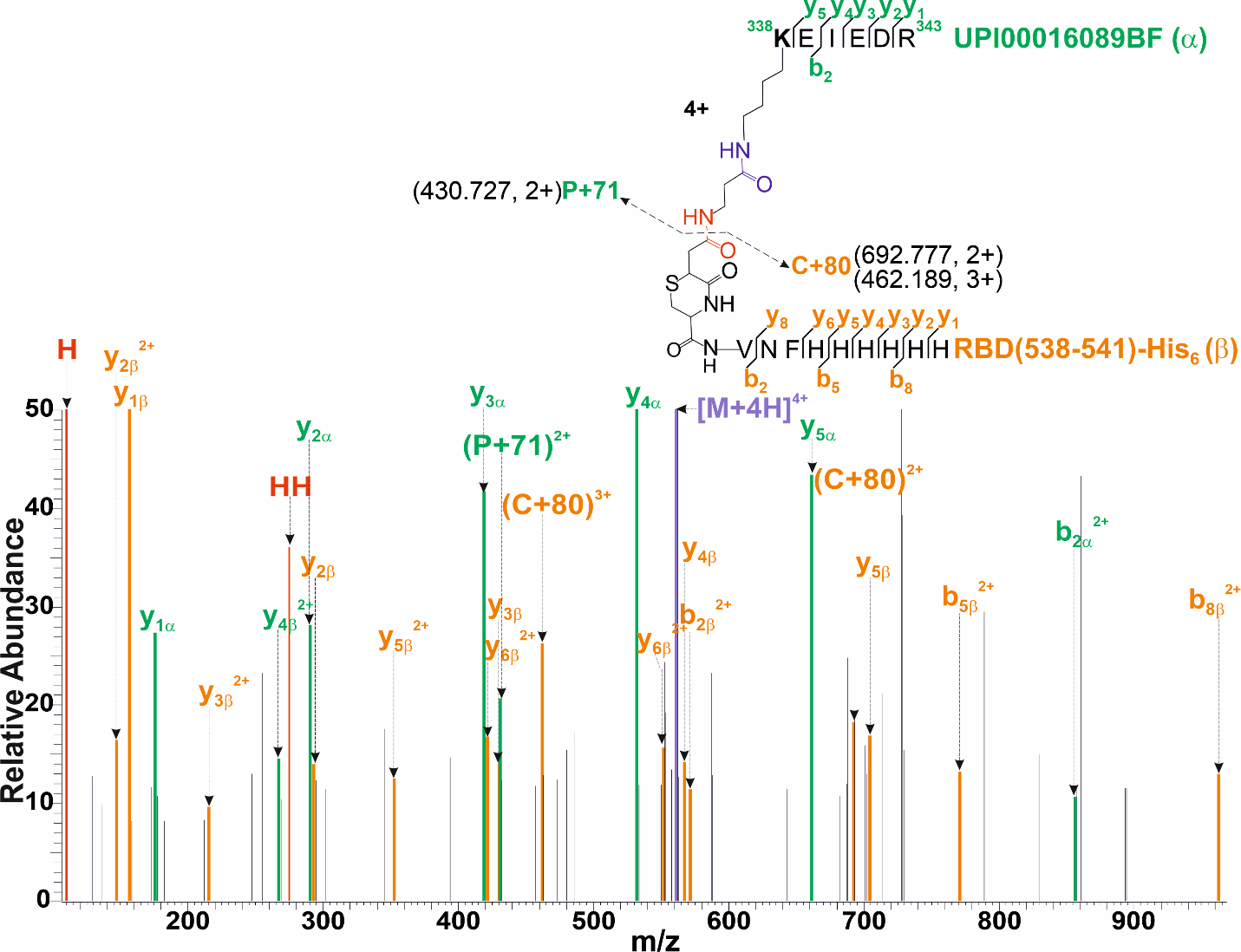

**Fig S21**. ESI-MS/MS spectrum assigned by pLink2 software [7] to a tryptic type 2 peptide composed by the *C*-terminal peptide of the RBD of SARS-CoV-2 (^538^**C**VNF^541^-HHHHHH) crosslinked via a hydrolyzed thiosuccinimide linker to the peptide [^31^DI**K**DEFVER^33^] of 3-hydroxybutyryl-CoA dehydrogenase (UPI00000106CD) from *C. tetani*. The (C+98) linker fragment ion [1, 9] detected at (*m/z* = 701.784, 2+ and *m/z* =1402.557, 1+), was generated by the fragmentation of the pseudopeptide bond (highlighted in red) newly formed due to the hydrolysis reaction [12, 13]. The signals at (*m/z* = 110.071, 1+ and *m/z* = 275.126, 1+) corresponds to the immonium ion of histidine (H) and to an internal fragment ion (HH) of the tandem repeat of six histidine residues present in type 2 tryptic peptides identified here, respectively.

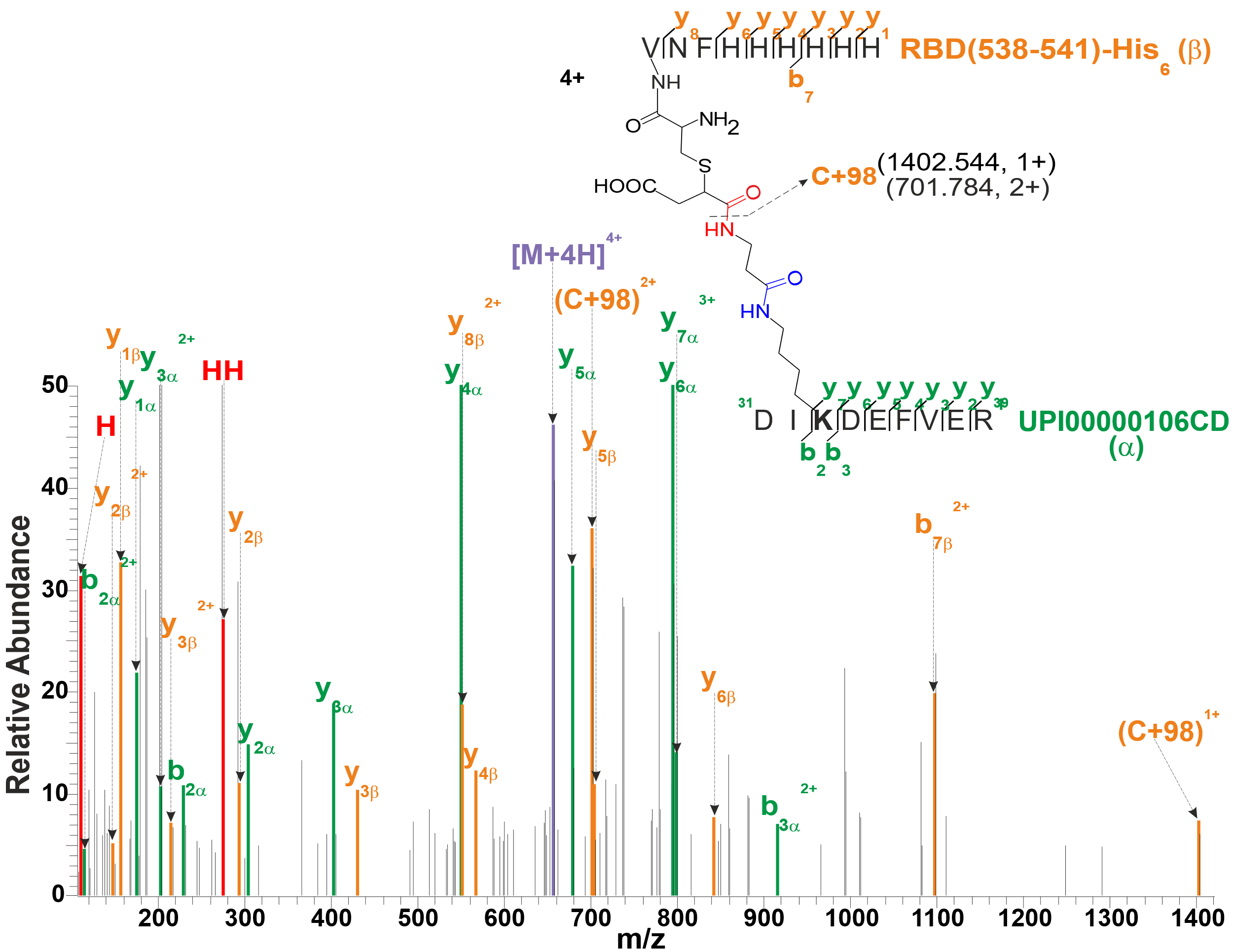
