## Supplemental Tables. for "LC-MS/MS characterization of SOBERANA^®^02, a receptor binding domain-tetanus toxoid conjugate vaccine against SARS-CoV-2"

**Affiliations:**

Avenida 31, e/ 158 y 190, Cubanacán, Playa. CP 11600, PO. Box 6162, La Habana 11600, Cuba.

**Index**

Table S1. Conjugation sites identification in ten proteins different to d-TeNT determined by LC-MS/MS analysis and based on the assignment of type 2 peptides with a transcyclized linker.

| # | Assignment ^i)^ | CS ^ii)^ | m/z theor ^iii)^ | m/z exp ^iii)^ | z | Error (ppm) | S/W ^iv)^ | P+71 (1), C+80 (2), H (3), HH (4), HHH (5) ^v)^ |
| --- | --- | --- | --- | --- | --- | --- | --- | --- |
| **(1) UPI0002DAC767 Putative S-layer protein/N-acetylmuramoyl-L-alanine amidase** | | | | | | | | |
| 1 | [^426^K-K^433^]-(^538^C-F^541^)-(His)_6_) | K426 | 575.778 | 575.775 | 4 | -5.387 | 1 | 1+2+4+5 |
| 2 | [^609^L-K^618^] -(^538^C-F^541^)-(His)_6_) | K613 | 515.436 | 515.433 | 5 | -6.741 | 1+2 | 1+2+4 |
|  |  |  | 644.043 | 644.040 | 4 | -5.049 | 1 | 2+4+5 |
| 3 | [^663^V-K^673^]-(^538^C-F^541^)-(His)_6_) | K669 | 536.652 | 536.649 | 5 | -6.482 | 1+2 | 1+2+3+4 |
|  |  |  | 670.563 | 670.560 | 4 | -4.828 | 1+2 | 2+3+4 |
|  |  |  | 894.082 | 894.079 | 3 | -3.100 | 1+2 | 4 |
| 4 | [^694^D-K^701^]-(^538^C-F^541^)-(His)_6_) | K698 | 583.272 | 583.269 | 4 | -5.949 | 1 | 1+2+4+5 |
| 5 | [^798^T-K^808^]-(^538^C-F^541^)-(His)_6_) | K803 | 549.252 | 549.252 | 5 | 0.000 | 2 | 1+2+4 |
| 6 | [^916^Y-K^927^]-(^538^C-F^541^)-(His)_6_) | K922 | 708.066 | 708.061 | 4 | -6.900 | 1 | 3+4 |
| 7 | [^928^D-K^936^]-(^538^C-F^541^)-(His)_6_) | K932 | 632.279 | 632.276 | 4 | -5.211 | 1 | 2+4 |
| 8 | [^984^T-K^992^]-(^538^C-F^541^)-(His)_6_) | K987 | 504.236 | 504.237 | 5 | 1.777 | 1 | 3+4+5 |
|  |  |  | 630.043 | 630.040 | 4 | -4.888 | 1+2 | 4 |
| 9 | [^1118^E-K^1126^]-(^538^C-F^541^)-(His)_6_) | K1125 | 492.819 | 492.816 | 5 | -6.047 | 1+2 | 1+2+4 |
|  |  |  | 616.023 | 616.023 | 4 | 0.000 | 1+2 | 2+3+4 |
| **(2) UPI00001A9618 Tyrosine phenol-lyase** | | | | | | | | |
| 1 | [^205^E-K^213^] -(^538^C-F^541^)-(His)_6_) | K209 | 409.534 | 409.530 | 6 | -8.665 | 1 | 2+4+5 |
|  |  |  | 491.039 | 491.036 | 5 | -6.993 | 1+2 | 1+2+4 |
|  |  |  | 613.547 | 613.544 | 4 | -4.307 | 1 | 2+4+5 |
| **(3) UPI00000106CD 3-hydroxybutyryl-CoA dehydrogenase** | | | | | | | | |
| 1 | [^31^D-R^39^]-(^538^C-F^541^)-(His)_6_) | K33 | 521.837 | 521.834 | 5 | -5.923 | 1+2 | 1+2+4 |
|  |  |  | 652.044 | 652.042 | 4 | -3.817 | 1+2 | 2+3+4+5 |
|  |  |  | 869.057 | 869.055 | 3 | -1.834 | 1+2 | 1+4+5 |
| 2 | [^276^G-K^282^]-(^538^C-F^541^)-(His)_6_) | K282 | 462.399 | 462.396 | 5 | -7.062 | 1 | 1+2+4+5 |
|  |  |  | 577.747 | 577.744 | 4 | -5.416 | 1 | 1+2+4+5 |
| **(4) UPI00016089BF Chaperonin GroEL** | | | | | | | | |
| 1 | [^338^K-R^343^]-(^538^C-F^541^)-(His)_6_) | K338 | 561.754 | 561.754 | 4 | 0.000 | 1+2 | 1+2+3+4 |
| 2 | [^345^V-R^360^]-(^538^C-F^541^)-(His)_6_) | K348 | 693.322 | 693.318 | 5 | -5.587 | 1+2 | 2 |
|  |  |  | 866.401 | 866.394 | 4 | -8.474 | 1+2 | 3+4 |
| 3 | [^361^E-R^366^]-(^538^C-F^541^)-(His)_6_) | K362 | 452.211 | 452.207 | 5 | -9.026 | 1+2 | 2+3+4 |
|  |  |  | 565.011 | 565.007 | 4 | -7.417 | 1+2 | 2+3+4 |
| 4 | [^379^V-R^390^]-(^538^C-F^541^)-(His)_6_) | K388 | 552.660 | 552.660 | 5 | 0.000 | 2 | 2+4+5 |
| **(5) UPI0000010752 Methylaspartate ammonia-lyase** | | | | | | | | |
| 1 | [^147^K-R^155^]-(^538^C-F^541^)-(His)_6_) | K147 | 498.440 | 498.437 | 5 | -6.904 | 1+2 | 2+4 |
|  |  |  | 622.798 | 622.795 | 4 | -4.643 | 1+2 | 2+4 |
| 2 | [^204^L-K^210^]-(^538^C-F^541^)-(His)_6_) | K207 | 550.767 | 550.764 | 4 | -4.564 | 1 | 1+2+4 |
| 3 | [^222^I-R^226^]-(^538^C-F^541^)-(His)_6_) | K224 | 420.416 | 420.413 | 5 | -7.443 | 1 | 1+2+4+5 |
|  |  |  | 525.018 | 525.015 | 4 | -6.351 | 1 | 2+4+5 |
| 4 | [^282^Q-R^289^]-(^538^C-F^541^)-(His)_6_) | K283 | 496.221 | 496.218 | 5 | -6.498 | 1+2 | 2+4 |
|  |  |  | 619.774 | 619.771 | 4 | -4.637 | 1 | 2+4+5 |
| **(6) UPI000001053D Cell wall-binding repeat-containing protein** | | | | | | | | |
| 1 | [^526^N-R^536^]-(^538^C-F^541^)-(His)_6_) | K533 | 654.303 | 654.303 | 4 | 0.000 | 2 | 4 |
| **(7) UPI0002DA4B64 Imidazolonepropionase** | | | | | | | | |
| 1 | [^20^K-K^28^] -(^538^C-F^541^)-(His)_6_) | K20 | 622.050 | 622.047 | 4 | -5.107 | 1 | 2+4+5 |
| 2 | [^159^N-K^169^]-(^538^C-F^541^)-(His)_6_) | K163 | 677.070 | 677.067 | 4 | -4.341 | 1 | 2+4 |
| 3 | [^159^N-K^169^]-(^538^C-F^541^)-(His)_6_) | K164 | 677.070 | 677.067 | 4 | -4.946 | 1 | 2+3+4+5 |
| 4 | [^164^K-R^171^]-(^538^C-F^541^)-(His)_6_) | K169 | 797.039 | 797.038 | 3 | -1.255 | 1 | - |
| **(8) UPI000001030F Aldehyde-alcohol dehydrogenase** | | | | | | | | |
| 1 | [^274^G-R^281^]-(^538^C-F^541^)-(His)_6_) | K279 | 478.019 | 478.016 | 5 | -6.672 | 1+2 | 2+4+5 |
|  |  |  | 597.272 | 597.269 | 4 | -5.533 | 1 | 2+4+5 |
| **(9) UPI000001056B Chaperone protein DnaK** | | | | | | | | |
| 1 | [^418^Q-R^428^]-(^538^C-F^541^)-(His)_6_) | K424 | 665.554 | 665.552 | 4 | -3.558 | 1 | 2+4+5 |
| **(10) UPI000001011B Protein with rubredoxin/rubrerythrin domain** | | | | | | | | |
| 1 | [^86^E-R^97^]-(^538^C-F^541^)-(His)_6_) | K96 | 564.264 | 564.264 | 5 | 0.000 | 2 | 2+4 |
|  |  |  | 705.078 | 705.077 | 4 | -1.418 | 2 | 2+4 |

i. Numbers inside brackets and parentheses correspond to the position of the type 2 tryptic peptides within the sequences of ten proteins of *C. tetani* and the C-terminal peptide of the recombinant RBD(R^319^-F^541^)-His_6_, respectively, crosslinked by a transcyclized linker.
ii. CS means conjugation sites. Conjugation sites identification. Indicates the position of several lysine residues in proteins of *C. tetani* that are linked by the transcyclized linker to the Cys^538^ of the RBD(R^319^-F^541^)-His_6_.
iii. The *m/z* theor and *m/z* exp correspond to the theoretical and experimental molecular masses of the type 2 tryptic peptides identified with the transcyclized linker.
iv. S/W means software. The numbers 1 and 2 mean that the MS/MS spectra were assigned by the pLink2 [24] and Kojak [25] software, respectively.
v. The numbers inside parentheses correspond to the diagnostic ions that were used in the validation process of the MS/MS spectra assigned to type 2 tryptic peptides. Linker fragment ions P+71, C+80, C-34, the immonium ion of H (*m/z* = 110.072) and internal ions HH (*m/z* = 275.126), HHH (*m/z* = 412.185) that were detected in the MS/MS spectra of the identified type 2 tryptic peptides are labeled with number 1, 2, 3, 4, and 5, respectively.

Table S2. Conjugation sites identification in seven proteins different to d-TeNT determined by LC-MS/MS analysis and based on the assignment of type 2 peptides with a hydrolyzed thiosuccinimide linker.

| # | Assigment ^i)^ | CS ^ii)^ | m/z theor ^iii)^ | m/z exp ^iii)^ | z | Error (ppm) | S/W ^iv)^ | P+71 (1), P+203 (2), C+98 (3), C-34 (4), H(5), HH (6), HHH (7) ^v)^ |
| --- | --- | --- | --- | --- | --- | --- | --- | --- |
| **(1) UPI0002DAC767 Putative S-layer protein/N-acetylmuramoyl-L-alanine amidase** | | | | | | | | |
| 1 | [^426^K-K^433^]-(^538^C-F^541^)-(His)_6_) | K426 | 580.281 | 580.28 | 4 | -5.698 | 1+2 | 6 |
| 2 | [^441^E-K^445^]-(^538^C-F^541^)-(His)_6_) | K443 | 697.334 | 697.328 | 3 | -8.808 | 1 | 2+6 |
| 3 | [^526^V-K^540^]-(^538^C-F^541^)-(His)_6_) | K533 | 649.302 | 649.299 | 5 | -4.594 | 1+2 | 2+3+6 |
|  |  |  | 811.375 | 811.374 | 4 | -1.324 | 1+2 | 2+3+6 |
| 4 | [^609^L-K^618^]-(^538^C-F^541^)-(His)_6_) | K613 | 519.036 | 519.035 | 5 | -2.460 | 1 | 1+3+6 |
|  |  |  | 648.543 | 648.54 | 4 | -8.510 | 1 | 3+5+6 |
| 5 | [^663^V-K^673^]-(^538^C-F^541^)-(His)_6_) | K669 | 540.252 | 540.250 | 5 | -2.824 | 1+2 | 1+3+6 |
|  |  |  | 675.313 | 675.313 | 4 | 0.043 | 1+2 | 3+6+7 |
| 6 | [^804^V-K^810^]-(^538^C-F^541^)-(His)_6_) | K808 | 800.700 | 800.695 | 3 | -6.603 | 1 | - |
| 7 | [^1046^I-K^1053^]-(^538^C-F^541^)-(His)_6_) | K1048 | 597.274 | 597.269 | 4 | -8.881 | 1 | 3+6+7 |
| 8 | [^1126^K-K^1130^]-(^538^C-F^541^)-(His)_6_) | K1126 | 428.203 | 428.202 | 5 | -3.014 | 1+2 | 6 |
|  |  |  | 535.002 | 535.002 | 4 | -0.323 | 1+2 | 6 |
| 9 | [^1141^F-K^1151^]-(^538^C-F^541^)-(His)_6_) | K1142 | 678.567 | 678.565 | 4 | -3.282 | 1 | 3+6 |
| **(2) UPI00000106CD 3-hydroxybutyryl-CoA dehydrogenase** | | | | | | | | |
| 1 | [^31^D-R^39^]-(^538^C-F^541^)-(His)_6_) | K33 | 525.435 | 525.435 | 5 | -0.798 | 1+2 | 3+5+6+7 |
|  |  |  | 656.798 | 656.795 | 4 | -4.472 | 1+2 | 3+5+6 |
|  |  |  | 875.060 | 875.059 | 3 | -1.677 | 1+2 | 1+3+6 |
| 2 | [^276^G-K^282^]-(^538^C-F^541^)-(His)_6_) | K282 | 466.001 | 465.998 | 5 | -6.278 | 1 | 1+3+6+7 |
|  |  |  | 582.250 | 582.247 | 4 | -5.483 | 1 | 3+6 |
| **(3) UPI0000010752 Methylaspartate ammonia-lyase** | | | | | | | | |
| 1 | [^147^K-R^155^]-(^538^C-F^541^)-(His)_6_) | K147 | 627.050 | 627.048 | 4 | -3.670 | 1+2 | 3+5+6 |
| **(4) UPI000001053D Cell wall-binding repeat-containing protein** | | | | | | | | |
| 1 | [^397^L-K^406^]-(^538^C-F^541^)-(His)_6_) | K402 | 860.066 | 860.060 | 3 | -6.976 | 1 | 6 |
| 2 | [^405^A-K^414^]-(^538^C-F^541^)-(His)_6_) | K406 | 870.413 | 870.414 | 3 | 1.277 | 1 | - |
| **(5) UPI0002DA4B64 Imidazolonepropionase** | | | | | | | | |
| 1 | [^20^K-K^28^]-(^538^C-F^541^)-(His)_6_) | K20 | 626.553 | 626.549 | 4 | -6.586 | 1+2 | 3+6+7 |
| **(6) UPI000001056B Chaperone protein DnaK** | | | | | | | | |
| 1 | [^418^Q-R^428^]-(^538^C-F^541^)-(His)_6_) | K424 | 536.247 | 536.243 | 5 | -6.549 | 1 | 3+6 |
| **(7) UPI000001011B Protein with rubredoxin/rubrerythrin domain** | | | | | | | | |
| 1 | [^168^H-K^173^]-(^538^C-F^541^)-(His)_6_) | K170 | 432.803 | 432.800 | 5 | -6.932 | 1 | 3+5+6+7 |

i. Type 2 peptides are composed by tryptic peptides of seven proteins of *C. tetani* crosslinked by a hydrolyzed thiosuccinimide linker to the *C*-terminal tryptic peptide ((^538^C-F^541^)-(His)_6_) of the recombinant RBD of SARS-CoV-2. Numbers inside brackets and parentheses correspond to the position of the type 2 tryptic peptides in the identified proteins and the recombinant RBD(R^319^-F^541^)-His_6_, respectively.
ii. Conjugation sites identification. Indicates the position of several lysine residues in proteins of *C. tetani* that are linked by a hydrolyzed thiosuccinimide linker to the Cys^538^ of the RBD(R^3 19^-F^541^)-His_6_.
iii. The *m/z* theor and *m/z* exp correspond to the theoretical and experimental molecular masses of the type 2 tryptic peptides identified with the hydrolyzed thiosuccinimide linker.
iv. The numbers 1 and 2 mean that the MS/MS spectra were assigned by the pLink2 [24] and Kojak [25] software, respectively.
v. The numbers inside parentheses correspond to the diagnostic ions that were detected in the validation process of the MS/MS spectra assigned to type 2 tryptic peptides with the hydrolyzed thiosuccinimide linker. Linker fragment ions P+71, P+203, C+98, and C-34, the immonium ion of His (*m/z* = 110.072) and internal ions HH (*m/z* = 275.126), and HHH (*m/z* = 412.185) are labeled with numbers 1, 2, 3 4, 5, 6 and 7, respectively.

Tabla S3. Identification of the conjugation sites by LC-MS/MS analysis of other low abundance proteins of *C. tetani* other than d-TeNT, which are also present in the TT preparation. The identification of the conjugation sites is based on the MS/MS assignment of linear peptides containing lysine residues cross-linked to the C-terminal peptides of the RBD by a transcyclized (K+1454.58) and hydrolyzed thiosuccinimide linker (K+1472.59).

| # | Assigment^v)^ | CS^vi)^ | m/z theo ^vii)^ | m/z exp ^vii)^ | z | Error  (ppm) | P+71 (1), P+203 (2), C+80 (3), C+98 (4),  C-34 (5), C+151 (6), C+169 (7), H+ (8)^viii)^ |
| --- | --- | --- | --- | --- | --- | --- | --- |
| **Linear peptides with the transcycled linker** | | | | | | | |
| (1)UPI0002DAC767 Trigger factor | | | | | | | |
| 1 | ^601^SL**K(+1454.58)**EC(+57.02)EIR^608^ | K(603) | 498.628 | 498.627 | 5 | -1.203 | 3+6 |
| 2 | ^609^LNED**K(+1454.58)**VSADK^618^ | K(613)* | 644.043 | 644.043 | 4 | -0.311 | 3+6 |
| 3 | ^663^VTFTAS**K(+1454.58)**DEVK^673^ | K(669)* | 670.563 | 670.562 | 4 | -1.640 | 3+6 |
| 4 | ^694^DVSE**K(+1454.58)**VAK^701^ | K(698)* | 583.272 | 583.271 | 4 | -1.029 | 3+6 |
| 5 | ^798^TSTTN**K(+1454.58)**VENYK^808^ | K(803)* | 548.848 | 548.851 | 5 | 5.102 | 1+3+6 |
|  |  |  | 685.562 | 685.563 | 4 | 0.729 | 1+3+6 |
| 6 | ^984^TIE**K(+1454.58)**EDLSK^992^ | K(987)* | 630.043 | 630.043 | 4 | 0.000 | - |
| (2) UPI00001A9618 Tyrosine phenol-lyase | | | | | | | |
| 1 | ^205^ELTA**K(+1454.58)**HGIK^213^ | K(203) | 491.039 | 491.038 | 5 | -1.833 | 1+3+6 |
| (3)UPI00000106CD 3-hydroxybutyryl-CoA dehydrogenase | | | | | | | |
| 1 | ^31^DI**K(+1454.58)**DEFVER^39^ | K(33)* | 521.837 | 521.837 | 5 | -0.192 | 1+3+6 |
|  |  |  | 652.044 | 652.044 | 4 | 0.460 | 3+6+8+9 |
| 2 | ^276^GFHDYS**K(+1454.58)**^282^ | K(282)* | 462.600 | 462.599 | 5 | -1.729 | 1+3+6 |
|  |  |  | 577.747 | 577.747 | 4 | -0.519 | 1+3+6 |
| (4) UPI00016089BF Chaperonin GroEL | | | | | | | |
| 1 | ^344^VNQI**K(+1454.58)**AQIEETTSEFDR^360^ | K(348)* | 693.523 | 693.524 | 5 | 1.009 | 3+6 |
|  |  |  | 866.401 | 866.401 | 4 | -0.577 | 6+8+9 |
| (5) UPI000001053D Cell wall-binding repeat-containing protein | | | | | | | |
| 1 | ^526^NEGAVTL**K(+1454.58)**DGR^536^ | K(533)* | 654.303 | 654.302 | 4 | -1.070 | - |
| (6) UPI000001030F Aldehyde-alcohol dehydrogenase | | | | | | | |
| 1 | ^274^GDEVD**K(+1454.58)**LR^281^ | K(279)* | 478.019 | 478.018 | 5 | -1.883 | 3+6 |
| **Linear peptides with the hydrolyzed linker** | | | | | | | |
| (1) UPI0002DAC767 Trigger factor | | | | | | | |
| 1 | ^526^VEYYLEE**K(+1472.59)**LPSGTNK^540^ | K(533)* | 649.303 | 649.303 | 5 | -0.154 | 1+2+4+7 |
| 2 | ^663^VTFTAS**K(+1472.59)**DEVK^673^ | K(669)* | 675.065 | 675.065 | 4 | 0.000 | 4 |
| (2) UPI00000106CD 3-hydroxybutyryl-CoA dehydrogenase | | | | | | | |
| 1 | ^33^DI**K(+1472.59)**DEFVER^39^ | K(33)* | 525.640 | 525.638 | 5 | -4.756 | 4+7+8 |
|  |  |  | 656.547 | 656.545 | 4 | -3.503 | 4+7+8 |
| (3) UPI000001053D Cell wall-binding repeat-containing protein | | | | | | | |
| 1 | ^526^NEGAVTL**K(+1472.59)**DGR^536^ | K(533) | 658.805 | 658.804 | 4 | -1.214 | 4 |
| (4) UPI0002DA4B64 Imidazolonepropionase | | | | | | | |
| 1 | ^20^**K(+1472.59)**DILIENGK^28^ | K(20)* | 626.548 | 626.549 | 4 | 0.798 | 4+7 |

i. Linear tryptic peptides within the sequences of seven proteins of *C. tetani*, where several lysines are modified with the linker, in its transcyclized and hydrolyzed forms, linked to the *C*-terminal tryptic peptide ((^538^C-F^541^)-(His)_6_) of the recombinant RBD of SARS-CoV-2. In peptide sequences, the letter in bold represent the modified lysine with the transcyclized (K(+1454.58)) and hydrolyzed linker (K(+1472.59)).
ii. CS means the conjugation sites. Conjugation sites identification. Indicate the position of several lysine residues in proteins of *C. tetani* that are linked by the transcyclized and hydrolyzed linker to the Cys^538^ of the RBD (^319^R-F^541^)-His_6_.
iii. The *m/z* theor and *m/z* exp correspond to the theoretical and experimental molecular masses of the linear tryptic peptides with the transcyclized and hydrolyzed linker, respectively.
iv. The numbers inside parentheses correspond to the diagnostic ions that were detected in the validation process of the MS/MS spectra assigned to linear tryptic peptides with the transcyclized and hydrolyzed linker, respectively. Linker fragment ions P+71, C+80 and C+151are in the case of the transcyclized linker, while P+71, P+203, C+98, C-34, C+169 represented the linker in the hydrolyzed form. The immonium ion of His (*m/z* = 110.072) is present in both forms of the linker. These are labeled with numbers 1, 2, 3 4, 5, 6, 7 and 8, respectively.

* Conjugation sites that identified by the both strategies used (Through linear peptides and type 2 peptides).
